# Synthetic peptide hydrogels as a model of the bone marrow niche demonstrate efficacy of a combined CRISPR-CAR T-cell therapy for acute myeloid leukaemia

**DOI:** 10.1101/2025.04.16.649170

**Authors:** W. Sebastian Doherty-Boyd, P. Monica Tsimbouri, Vineetha Jayawarna, Matthew Walker, Aqeel F. Taqi, Niall Mahon, Dominic Meek, Peter Young, Aline Miller, Adam West, Manuel Salmeron-Sanchez, Matthew J. Dalby, Hannah Donnelly

**Affiliations:** Centre for the Cellular Microenvironment, Advanced Research Centre, University of Glasgow, Glasgow, G12 8QQ, United Kingdom; Epigenetics Research Unit, School of Cancer Sciences, Wolfson Wohl Cancer Research Centre, University of Glasgow, Glasgow, G61 1QH, United Kingdom; Dasman Diabetes Institute, 15462, Dasman, Kuwait; School of Chemical Engineering and Analytical Sciences, University of Manchester Oxford Road, Manchester, M13 9P

## Abstract

Leukaemias, driven by mutations in hematopoietic stem cells (HSCs), rely on interactions with the bone marrow (BM) niche and other cell populations such as mesenchymal stromal cells (MSCs) for growth and survival. While chimeric antigen receptor (CAR) T-cell therapy shows promise for other hematological malignancies, its application to acute myeloid leukaemia (AML) is hindered by tumour heterogeneity and off-target toxicity. Combining CRISPR-Cas9 gene editing with CAR T-cell therapy has potential for selectively targeting AML cells while sparing healthy tissue. However, validating the efficacy of these treatments prior to clinical trial is hampered by the differences between humans and animal models typically used for pre-clinical testing. Furthermore, traditional in vitro models fail to replicate the complexity of the BM niche and often overestimate treatments’ efficacy. Here, we present a bioengineered human-cell containing BM niche model combining a fibronectin-presenting polymeric surface and a synthetic peptide hydrogel (PeptiGel) that mimics native BM tissue’s mechanical properties. This platform supports niche phenotypes in MSCs and HSCs and enables the evaluation of combined CRISPR-CAR T-cell therapy, demonstrating potential as a preclinical human model for testing novel therapies.

## Introduction

Leukaemias are a form of haematological malignancies characterised by the abnormal production of blood cells. Leukemic malignancy is commonly driven by mutated hematopoietic stem cells (HSCs), which become chemoresistant leukaemia stem cells (LSCs) and represent a major contributor to disease progression and relapse [1]. Healthy HSCs rely on local interactions with their microenvironment, including other cell types such as mesenchymal stromal cells (MSCs) and the extracellular matrix (ECM) of the bone marrow where they reside. This specialised microenvironment is termed the bone marrow niche [2]. Likewise, LSCs depend on the BM niche; during leukemic transformation the niche is remodelled to facilitate LSC growth and survival, and has been shown to play a role in chemoresistance [3]. The niche should therefore be considered during therapeutic development.

Adoptive chimeric antigen receptor (CAR) T-cell transfer has emerged as a promising treatment for relapsed and refractory haematological malignancies [4–6], with potentially curative therapies for B cell malignancies reaching FDA approval [7,8]. However, progress in developing CAR T-cell therapies for acute myeloid leukaemia (AML), an aggressive blood cancer characterised by immature myeloid lineage cells, remains limited [9]. AML, the most common leukaemia in adults, exhibits significant heterogeneity and intrinsic tumour variability, complicating prediction of patient responses to treatment and highlighting the need for novel therapeutic approaches [1]. Many in vitro and in vivo studies have demonstrated the potential of CAR T-cells targeting surface proteins such as CD33 [6], CD123 [10], and CLL-1 [11] to ablate AML cells. However, clinical trials have shown low response rates accompanied by ‘on-target off-tumour’ toxicity due to the frequent expression of the target on healthy haematopoietic cells [12]. An approach combining CRISPR-Cas9 gene editing with CAR T-cell therapy has shown potential to increase the resistance of healthy cells by reducing or eliminating their expression of the target antigen [13]. In AML, it was demonstrated that using this method to target the myeloid cell surface marker CD33, which is expressed on 85-90% of AML patients’ cells [14–17], led to selective ablation of AML cells while sparing healthy cells [18]. However, to our knowledge, this CRISPR-CAR T-cell approach has yet to be tested beyond animal models.

Currently, preclinical testing of CAR T-cell therapies relies on simplified in vitro systems, and in vivo models that fail to sufficiently predict efficacy or adverse events that occur in patients, demonstrating a significant translational gap [19–21]. Traditional in vitro assays are often 2-dimensional (2D) cultures or co-cultures used in early-stage development to assess cancer cell-killing ability that fail to consider off-target effects on healthy cells or interactions with the tumour microenvironment [22]. Furthermore, animal models used are often immunodeficient or have significant difference to human immunity and often overestimate the efficacy of the treatment [23,24]. This highlights the need for more human-relevant models from the earliest stages of immunotherapy development [22,25].

The development of bioengineered humanised (human cell containing) in vitro platforms is showing promise for modelling and testing therapeutic approaches. BM models range from simple liquid culture to highly complex hydrogel-based co-culture and organoid systems that faithfully recapitulate the 3D complexity of the BM microenvironment [26]. However, many 3D systems rely on animal derived scaffolds or hydrogels derived from collagens or Matrigel, leading to significant batch-to-batch variability, reducing the physiological relevance of these systems and the reproducibility required for therapeutic testing [22].

In this study, we present the development of a bioengineered model of the BM niche and demonstrate its application for investigating the efficacy of a combined CRISPR-CAR T-cell therapy. We employ a polymeric surface that promotes the formation of physiological-like networks of fibronectin (FN), a BM-abundant ECM protein, thereby enhancing the attachment of growth factors (GFs) and supporting the adhesion of supporting MSCs [27–31]. We then incorporate a synthetic, self-assembling peptide hydrogel (Peptigel), with tuneable mechanical properties, which eliminates all non-serum animal components [32,33]. Our previous work has shown that low-stiffness ECM supports stromal stem cell phenotypes typical of the BM niche, which, in turn, support clinically relevant HSCs in vitro [34]. Here, the platform replicates the mechanical complexity of the endogenous BM niche, providing a supportive environment for HSCs and facilitating the evaluation of combined CRISPR-CAR T-cell therapy efficacy.

## Results

### Material characterisation

To mimic properties of the BM microenvironment, we designed a 3D, multicellular niche with mechanical and chemical elements akin to the BM niche (Fig. 1A). Our previous work has demonstrated that the polymer poly (ethyl acrylate) (PEA) can drive unfolding of the bone marrow-abundant ECM molecule FN [35], causing it to adopt a biomimetic, fibrillar network conformation [36]. Usually a cell-driven process, this exposes key domains, namely a heparin (P5F3) binding domain that allows GFs to be sequestered and presented, and a cell adherent RGD domain [37]. This has been applied for bone regeneration and MSC phenotype control in vitro [30] and in vivo [31] and recently in a BM niche model able to maintain HSCs in vitro [38]. As such we first PEA coated polymer cover slip (PCS) plates using plasma polymerisation of ethyl acrylate monomer. PCS plates were selected as the bottom of these plates consisted of a thin cover slip made of a poly(ethylene) derivative which facilitated increased microscopy image quality. Incorporation of PEA onto the surface was assessed with X-ray photoelectron spectroscopy (XPS) analysis (Fig. 1B). A significant O1 peak representative of PEA coating was detected on our PEA coated samples but not on standard uncoated controls (PCS). Furthermore, the spectra demonstrated similarity to previously reported PEA spectra [31,39,40].

**Figure 1.**
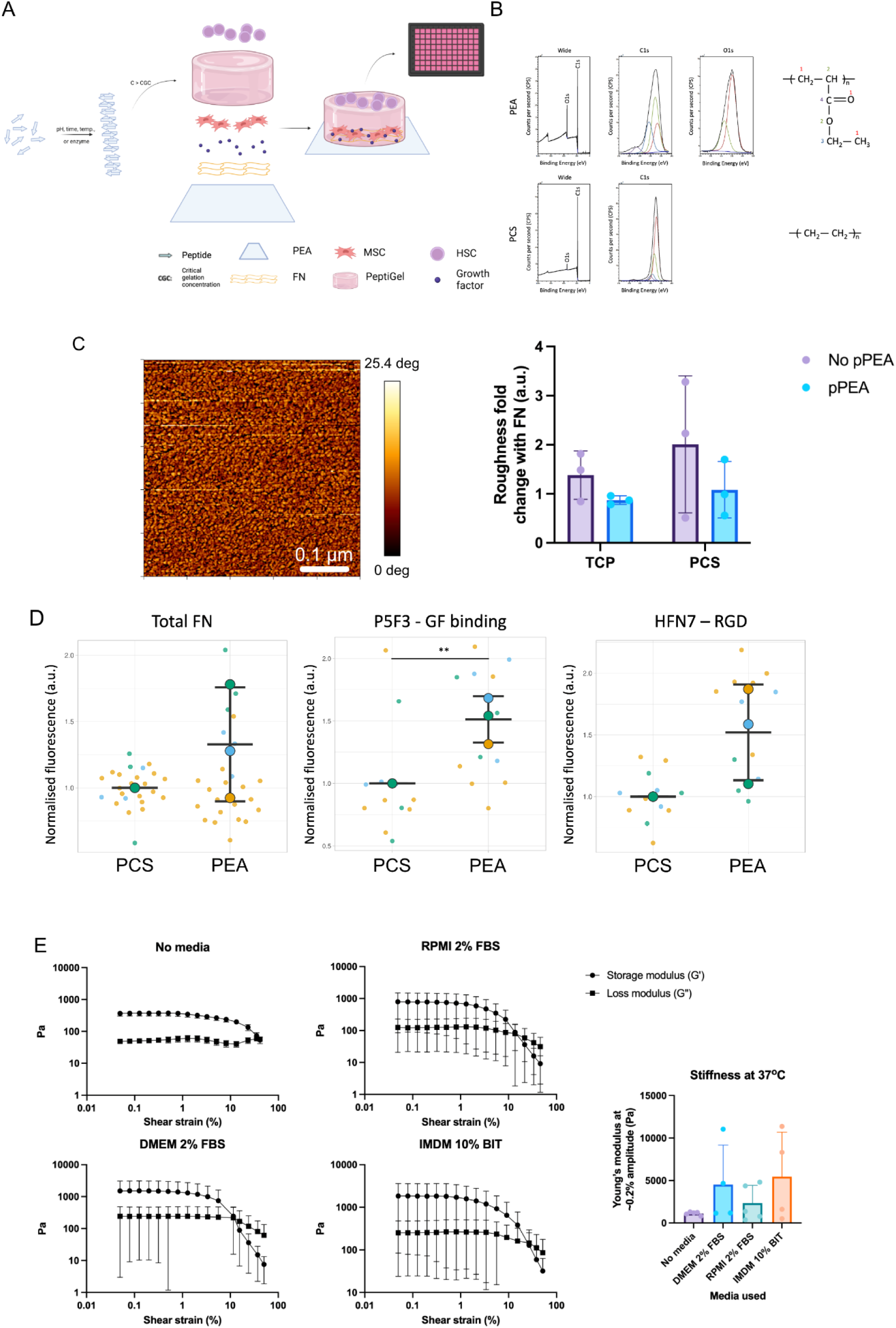
Characterising mechanical and chemical properties of a synthetic BM niche model. **A.** Schematic of the synthetic BM model. Created with BioRender.com. **B.** XPS spectra of polymer cover slip plates that were either PEA coated (PEA) or uncoated (PCS). From left to right: wide field, C1s, O1s spectra, chemical structures of surfaces; PEA and poly(ethylene), a derivative of which PCS surfaces were made of. **C.** Representative AFM images of PEA with FN, as well as fold change in roughness (Sa) of PEA and PCS surfaces when FN is included. **D** Total FN absorbed and FN domain availability on PCS and PEA surfaces, quantified using in-cell western analysis, normalised against total FN**. E** Rheological analysis by amplitude sweeps of PeptiGel incubated overnight with relevant cell culture medias. Graphs show mean ± SD. Statistics for **D** by unpaired, two-tailed t-test. *p < 0.05, **p < 0.01, non-significant not shown. **C-E** n=3 experimental replicates, **F** n=4 or 5 experimental replicates.

Next, FN was adsorbed onto PEA and PCS surfaces and formation of interconnected networks was confirmed only on PEA using atomic force microscopy (AFM) (Fig. 1C). To assess FN unfolding and exposure of key domains, fluorescent detection of antibodies that detect full length FN, the RGD domain and the P5F3 GF binding site were used. Fluorescent detection then demonstrated no significant difference, but a trend toward an increase in total amount of FN adsorbed and exposure of the RGD domain, while a significant increase in the availability of the P5F3 GF binding site on PEA compared to control PCS surfaces was observed (Fig. 1D), suggesting an unfolded FN conformation. Then, the osteogenic GF bone morphogenetic protein 2 (BMP2), which has previously been shown to stimulate a niche-like phenotype in MSCs in similar models [38], was adsorbed onto FN, facilitating synergistic, solid-phase GF presentation [30,41].

The BM niche is a low-stiffness hydrogel [42,43]. To create a 3D environment that was mechanically similar to endogenous BM, peptide-based hydrogels, termed PeptiGel, were used. PeptiGels are short amphipathic peptides (FEFKFEFK) assembled into B-sheet fibrils that form a 3D liquid retentive mesh [33]. The mechanical properties of PeptiGel when incubated with MSC media (2% DMEM), AML cell media (10% RPMI) and HSC media (IMDM+) was tested via rheology, as different media has been shown to affect gel stiffness [44] (Fig. 1E). The storage modulus was used to calculate gel stiffness. All media conditions caused PeptiGel stiffness to increase from ∼1000 Pa in the no media controls to ∼4500 Pa in 2% DMEM, ∼2300 Pa in 2% RPMI, and ∼5500 Pa in IMDM+. This may have been due to the media changing the gel’s pH, or charge screening [45]. Whilst a high degree of variance was observed, the average stiffness values of PeptiGel with all media types fell within the reported 1-10^4^ Pa range of soft tissues like BM, particularly the stiffer endosteal region [46–48].

In addition, we hypothesized that gel dilution would alter PeptiGel mechanical properties by changing peptide concentration. PeptiGel temperature can also affect mechanical properties by altering the kinetic energy of the gel components. Therefore, the effects of gel dilution and temperature on gel stiffness and viscoelasticity, measured as tan delta, as well as shear thinning properties, were assessed without media. Collagen type I gel, a natural animal-derived system that has previously been shown to produce a niche-like microenvironment in MSCs [34] was used as a control (Supplementary Fig. 1A-B). The temperatures tested were 37°C (incubation temperature) and 20°C (handling temperature). At 20°C, decreasing PeptiGel concentrations from 30 - 1 mg/ml caused a significant decrease in stiffness from ∼1300 Pa to ∼400 Pa across dilutions tested. At 37°C, a similar trend of decreased gel concentration leading to decreased stiffness (30 mg/mL ∼1100 Pa - 5 mg/mL gel ∼550 Pa) was observed. Gel viscoelasticity was also shown to be affected by gel dilution, where lower concentrations caused viscous properties to dominate at lower shear strain, represented by an increase in tan delta. Dilution increased gel fragility, demonstrated by transition to a tan delta greater than 1 at a lower shear strain, indicating that the gels were losing structural integrity (Supplementary Fig. 1C). The effect of media on PeptiGel’s viscoelasticity was shown to be minimal (Supplementary Fig. 1D). 30 mg/mL PeptiGel was also found to have sheer thinning properties, allowing easy deposition into niche models (Supplementary Fig. 1E). PeptiGel was significantly stiffer than collagen type I hydrogels, which may replicate the softer central regions of the BM niche [48,49] as opposed to the stiffer regions of the niche closer to the bone surface possibly replicated by the PeptiGel-based system [38].

### Model niche promotes a bone marrow niche phenotype in MSCs and HSCs

MSCs have been used as feeder layers to facilitate HSC retention in BM models for decades [26]. Viability of MSCs cultured under fully synthetic Peptigel (30mg/mL) in our PEA-FN-BMP2 niche model was expectedly slightly decreased compared to models which were 2D or included natural collagen type I hydrogel controls. However, live cells had similar morphology to these controls (Supplementary Fig. 2). MSCs with niche-like phenotypes are hypothesised to provide paracrine signals similar to MSCs in the native BM niche and therefore support HSCs in vitro. Nestin, an intermediate filament protein, and stem cell factor (SCF), a HSC regulatory cytokine, are highly expressed by subsets of HSC-regulating MSCs in the bone marrow microenvironment [50–54]. Nestin expression has also been demonstrated to be regulated by microenvironment mechanics [38,55]. The effect of PeptiGels with different peptide concentrations ranging from 1 to 30 mg/mL on MSC expression of nestin and SCF in our model BM niche was tested by immunofluorescence (IF) microscopy. 30 mg/mL PeptiGel led to the highest level of nestin and SCF expression (Fig. 2A-B, Supplementary Fig. 3B), resulting in a 1.6- and 1.5-fold increase respectively compared to 1 mg/mL PeptiGel. Interestingly, the expression of CXC chemokine ligand 12 (CXCL12), another HSC regulatory cytokine associated with egression from the BM [56], was also assessed and showed a 2.5-fold decrease in 30 mg/mL PeptiGel compared to 1mg/mL gel (Supplementary Fig. 3). Therefore, 30 mg/mL Peptigel was selected as the optimum formulation and used in the following BM model.

**Figure 2.**
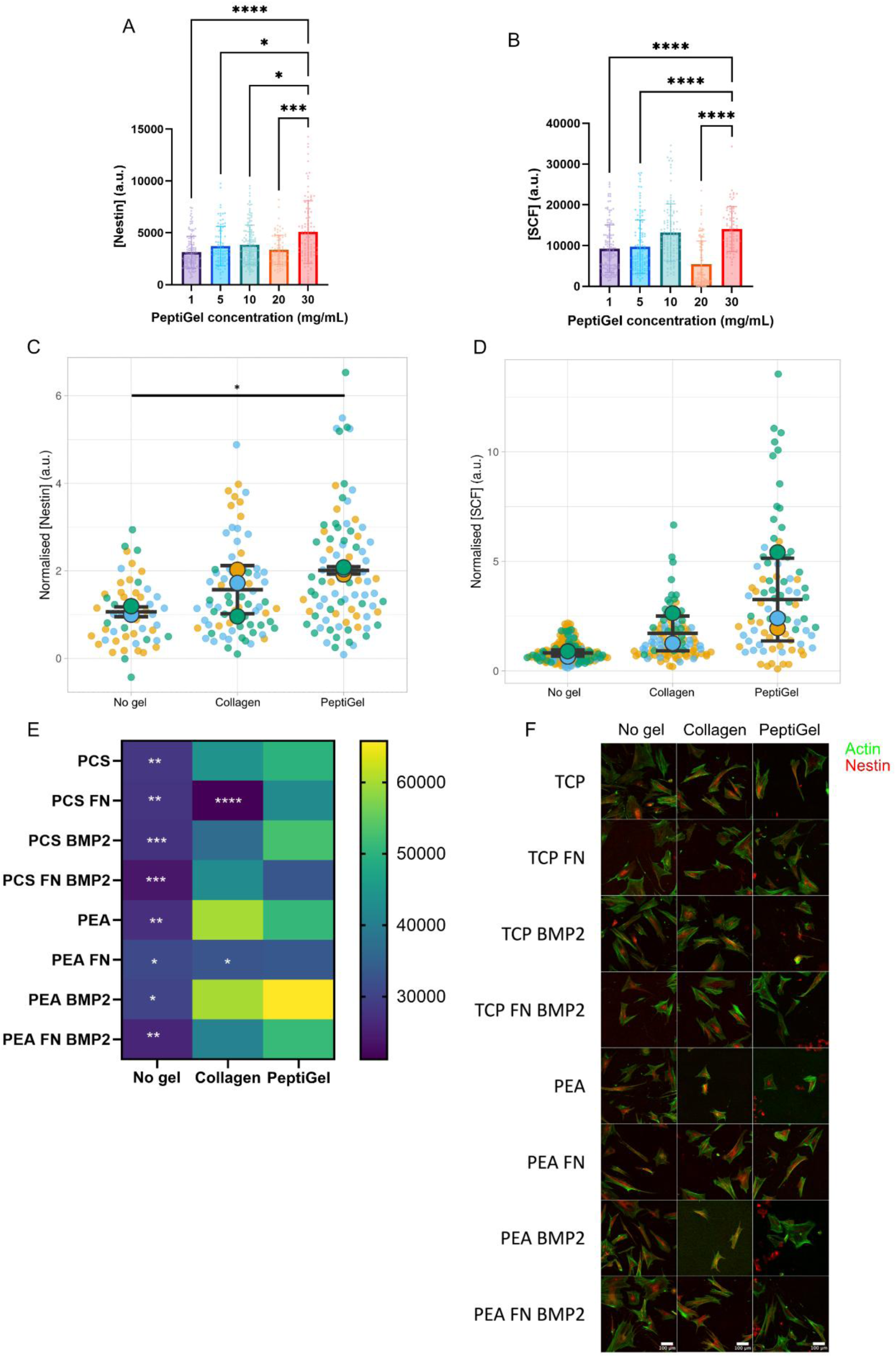
Phenotypic analysis of MSCs in a synthetic BM niche. Effect of gel dilution on **A** nestin and **B** SCF assessed by IF microscopy; representative images shown in supplementary fig. 5. **C** Nestin expression of MSCs grown in PeptiGel niches, as well as collagen and no gel niche controls, assessed by IF microscopy. **D** SCF expression in niches, assessed by IF microscopy. **E** Nestin expression in various environments including with or without PEA, FN and BMP2 coatings, PeptiGel, collagen and no gel conditions, assessed by IF microscopy. **F** Representative IF microscopy images of MSCs in various conditions. Scale bars are 100 µm, green = actin, red = nestin. Graphs show mean ± SD, heat map shows mean. Statistics for **A-C** by one-way ANOVA followed by Tukey multiple comparison test, for **E** by two-way ANOVA followed by Tukey multiple comparison test. Significance of each condition against PEA-FN-BMP2 +PeptiGel condition shown for **E**. *p < 0.05, **p < 0.01, ***p < 0.001 and ****p < 0.0001 non-significant not shown. **C** and **D** n=3 biological replicates, each consisting of 3 experimental replicates of ∼10 cells, **E** n=3 experimental replicates of ∼10 cells.

Next, nestin expression in PEA-FN-BMP2 +PeptiGel niches was compared against PEA-FN-BMP2 +collagen and PEA-FN-BMP2 no gel controls using IF microscopy (Fig. 2C-D). Here we show Peptigel-based niches support increased levels of nestin and SCF. This was confirmed with western blot and qPCR (Supplementary Fig. 4). The effect of each niche system ±PEA/FN/BMP2 on nestin expression was also tested (Fig. 2E-F), with the PEA-FN-BMP2 +PeptiGel system causing increased nestin expression. CXCL12 expression was assessed but found to have variable expression in the Peptigel system and no significant differences when compared to no gel control of the collagen gel system (Supplementary Fig. 5). Furthermore, MSCs in niche systems were cultured in mulitple cell culture medias required for co-cultures to investigate effect on stromal cell phenotype, assessed via nestin expression (Supplementary Fig. 6). However, no effect was observed between different media formulations.

Once the optimal niche for promoting the desired nestin^+^ SCF^+^ MSC phenotype was established, HSCs were introduced. HSCs were added to the PEA-FN-BMP2 ±gel niche systems after 7 days MSC culture, and co-cultured in the system for 5 days. Support of HSC ex vivo culture was investigated using flow cytometry (Fig. 3A-E). When investigating HSC phenotype, MSCs were found to increase haematopoietic cell (CD45^+^), hematopoietic stem and progenitor cell (HSPC) (CD45^+^ CD34^+^ CD38^-^) and long-term (LT-)HSC (CD45^+^ CD34^+^ CD38^-^ CD90^+^ CD45RA^-^) [57,58] numbers when included in no gel, collagen and PeptiGel systems. Notably, the LT-HSC population had the greatest fold change increase in the PeptiGel system, suggesting MSCs in this system were able to better support this clinically relevant population (Fig. 3E). HSC differentiation [59] in the three niche systems (PEA-FN-BMP2 +no gel/collagen/peptigel) ±MSCs was also measured (Fig. 3F-G). The inclusion of MSCs caused an increase in T-cell count, while the PeptiGel system resulted in an increased megakaryocyte count.

**Figure 3.**
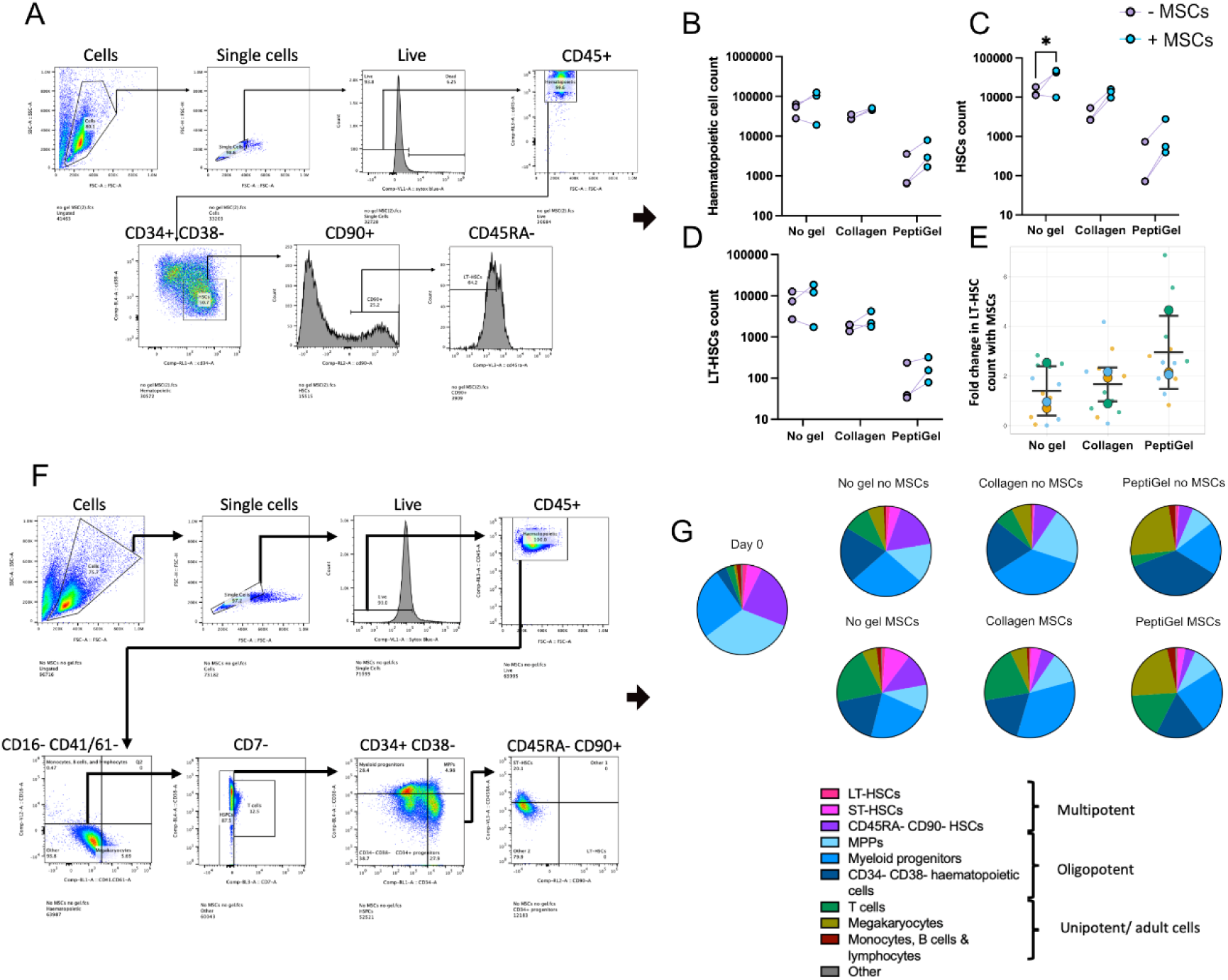
Phenotypic analysis of HSCs in synthetic BM niches. **A** Flow gating strategy for haematopoietic cell characterisation used for flow cytometry. **B** Effect of MSC inclusion on haematopoietic cell (live CD45+ cells) count after 5 days in PEA-FN-BMP2 no gel, +collagen gel, +Peptigel niche systems, assessed by flow cytometry. **C** HSC (live CD45+ CD34+ CD38-) count and **D** LT-HSC (live CD45+ CD34+ CD38- CD90+ CD45RA-) [57,58] counts were similarly assessed. **E** Fold change in LT-HSC count when MSCs were included in model systems. **F** Flow gating strategy for haematopoietic cell differentiation characterisation used for flow cytometry analysis. **G** Total haematopoietic cell population assessed by flow cytometry after 5 days in model BM niches. Cell types were defined as: Megakaryocytes (CD45+ CD16- CD41/61+), monocytes, B cells and lymphocytes (CD45+ CD16+ CD41/61-), T-cells (CD45+ CD16- CD41/61- CD7+), myeloid progenitors (CD45+ CD16- CD41/61- CD7- CD34- CD38+), CD34- CD38- haematopoietic cells (CD45+ CD16- CD41/61- CD7- CD34- CD38-), multipotent progenitors (CD45+ CD16- CD41/61- CD7- CD34+ CD38+), ST-HSCs (CD45+ CD16- CD41/61- CD7- CD34+ CD38- CD45RA+ CD90-), LT-HSCs (CD45+ CD16- CD41/61- CD7- CD34+ CD38- CD45RA- CD90+), CD45RA- CD90- HSCs (CD45+ CD16- CD41/61- CD7- CD34+ CD38- CD45RA- CD90-) [59]. Graphs show mean ± SD. **B-E** n=3 biological replicates, each consisting of 4 experimental replicates. **G** n=3 biological replicates, each consisting of 4 experimental replicates, except for day 0 which was n=3 biological replicates, each consisting of 3 experimental replicates.

### Genetic inactivation of CD33 in HSCs

We next sought to genetically edit healthy HSCs to remove the CAR T-cell target CD33. We hypothesized that using CRISPR-Cas9 approach with multiple single guide RNAs (sgRNA) that target different regions of the target gene would improve efficiency [60,61]. Therefore, to efficiently disrupt CD33 in healthy haematopoietic cells, we designed a CRISPR-Cas9 system using a cocktail of sgRNAs [62] to target three sites on the CD33 gene. These sites were in exons 2 and 3 of the gene, and in a putative upstream promoter (Fig. 4A). We first optimised the system in an AML cell line with high levels of CD33 expression (THP-1s) (Supplementary Fig. 7), then tested its efficiency in primary human HSCs at a DNA level with tracking of indels by decomposition (TIDE) analysis, which assesses DNA sequence alignment (Fig. 4B-D). At all sites tested, the sgRNA cocktail caused increased editing efficiency compared to each sgRNA individually. In addition, TIDE analysis was used to characterise the edits (Supplementary Fig. 8) and demonstrate short insertion/deletion (indel) mutations were occurring. HSCs do not express CD33; it is expressed in high levels on the surface of their progeny, differentiated myeloid cells [16]. Therefore, to investigate CD33-KO at the protein level, edited HSCs were differentiated down the myeloid lineage for 9 days. This was confirmed by increased expression of the myeloid marker CD13 and lineage cocktail staining across all conditions, demonstrating that HSC differentiation occurred and was not affected by CD33 knock-out (Fig. 4F-G). Using flow cytometry to detect surface level CD33, we found CD33 expression increased over time in unedited control HSCs, but not in edited HSCs, implying that the CRISPR-Cas9 knock-out prevented accumulation of CD33 on the edited HSCs’ (henceforth referred to as CD33^del^ HSCs) surface as they differentiated (Fig. 4H-J).

**Figure 4.**
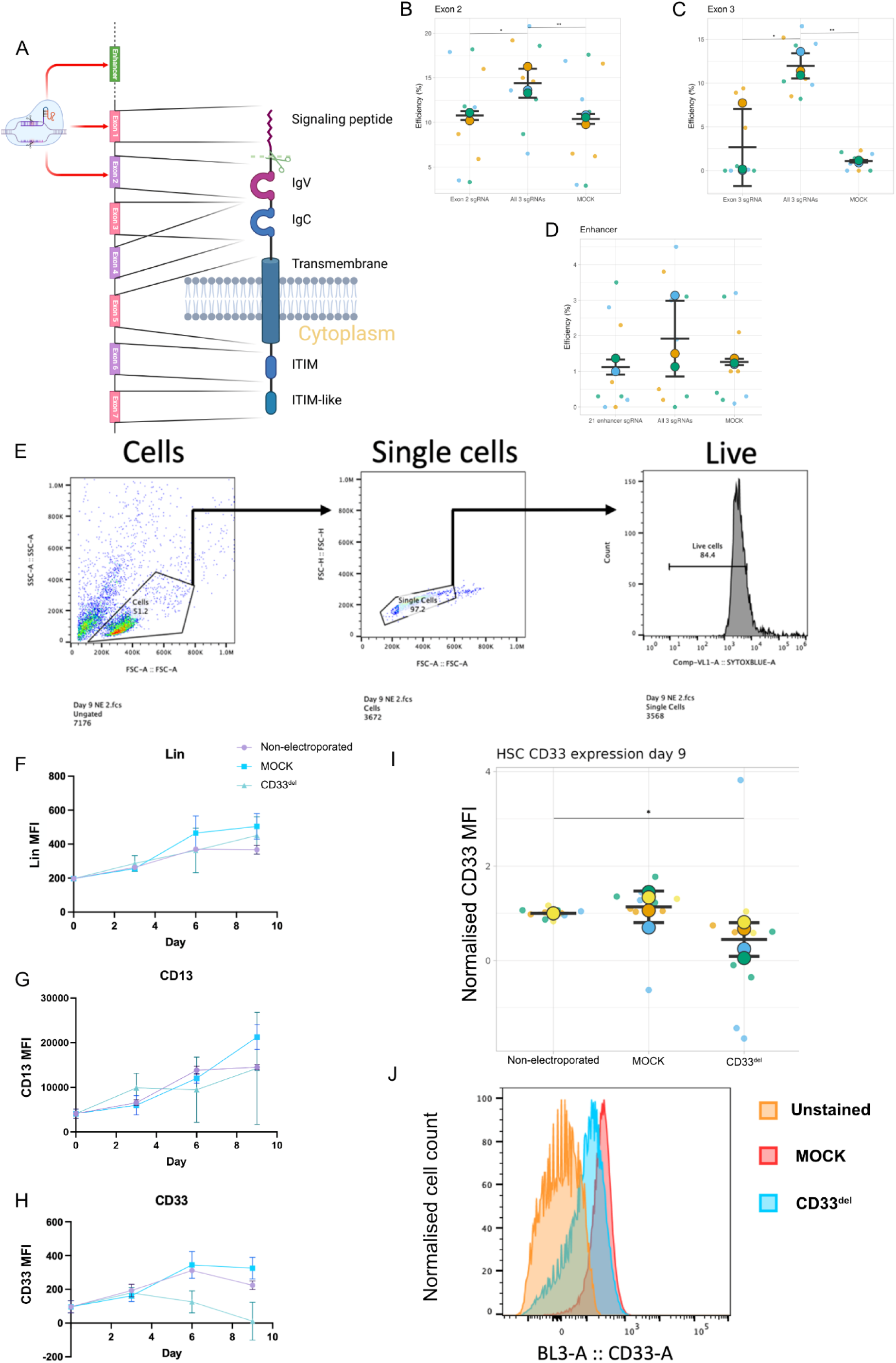
Genetic disruption of CD33 in HSCs. **A** Schematic of the CD33 gene with corresponding domains and sgRNA target sites. Created with BioRender.com. **B-D** Efficiency of gene knock-out at sgRNA target sites in B exon 2, **C** exon 3 and **D** an upstream enhancer of CD33 when individual sgRNAs, a cocktail of sgRNAs targeting all three sites or no sgRNAs (MOCK) were used for CRISPR, assessed with TIDE analysis. **E** Flow gating strategy for determining CD33 expression in HSCs. **F-H** Median fluorescence intensity (MFI) of HSCs stained with **F** lineage cocktail markers, **G** CD13, and **H** CD33 antibodies in corresponding channels over 9 days, assessed with flow cytometry. **I** Normalised CD33 expression in HSCs 9 days after editing, assessed by flow cytometry. **J** Representative flow cytometry fluorescence histogram of HSCs that were unstained, MOCK treated and edited with a cocktail of all three sgRNAs (CD33^del^). Graphs show mean ± SD. Statistics for **B-C** by one-way ANOVA followed by Tukey multiple comparison test, for **I** by Brown-Forsyth and Welch one-way ANOVA followed by Dunnett T3 multiple comparisons test. *p < 0.05, **p < 0.01, ***p < 0.001 and ****p < 0.0001, non-significant not shown. **B-D** n=3 experimental replicates, each consisting of 3 technical replicates, **F-H** n=3 experimental replicates, **I** n=4 biological replicates, each consisting of 2 or 3 experimental replicates.

### CD33-redirected CAR T-cells selectively ablate CD33+ cells

To test the effectiveness of CD33-redirected CAR T-cell on unedited and CD33^del^ cells, CAR T-cells were initially co-cultured with THP-1s (Fig. 5A) and cell death analysed using flow cytometry to determine the optimum target:effector (T:E) ratio (Fig. 5A). A 1:1 ratio was found to be sufficient for complete ablation of unedited THP-1 cells, while causing no significant drop in CD33^del^ THP-1 cell count (Fig. 5C-D). CAR T-cells effect on CD33^del^ and unedited HSCs was then assessed. Unedited cell numbers were shown to decrease in the presence of CD33-redirected CAR T-cells, while CD33^del^ HSC numbers remained high (Fig. 5E), suggesting that the CD33 edit was providing resistance to the CD33-redirected CAR T-cells. CAR T-cell addition was also shown to decrease CD33 expression across conditions (Fig. 5F). Since this drop in HSCs’ surface CD33 levels wasn’t accompanied by a drop in cell number in edited HSCs, other mechanisms besides CD33 presenting cells’ ablation, such as antigen escape [63], could have caused the drop in CD33 seen in response to the CAR T-cells.

**Figure 5.**
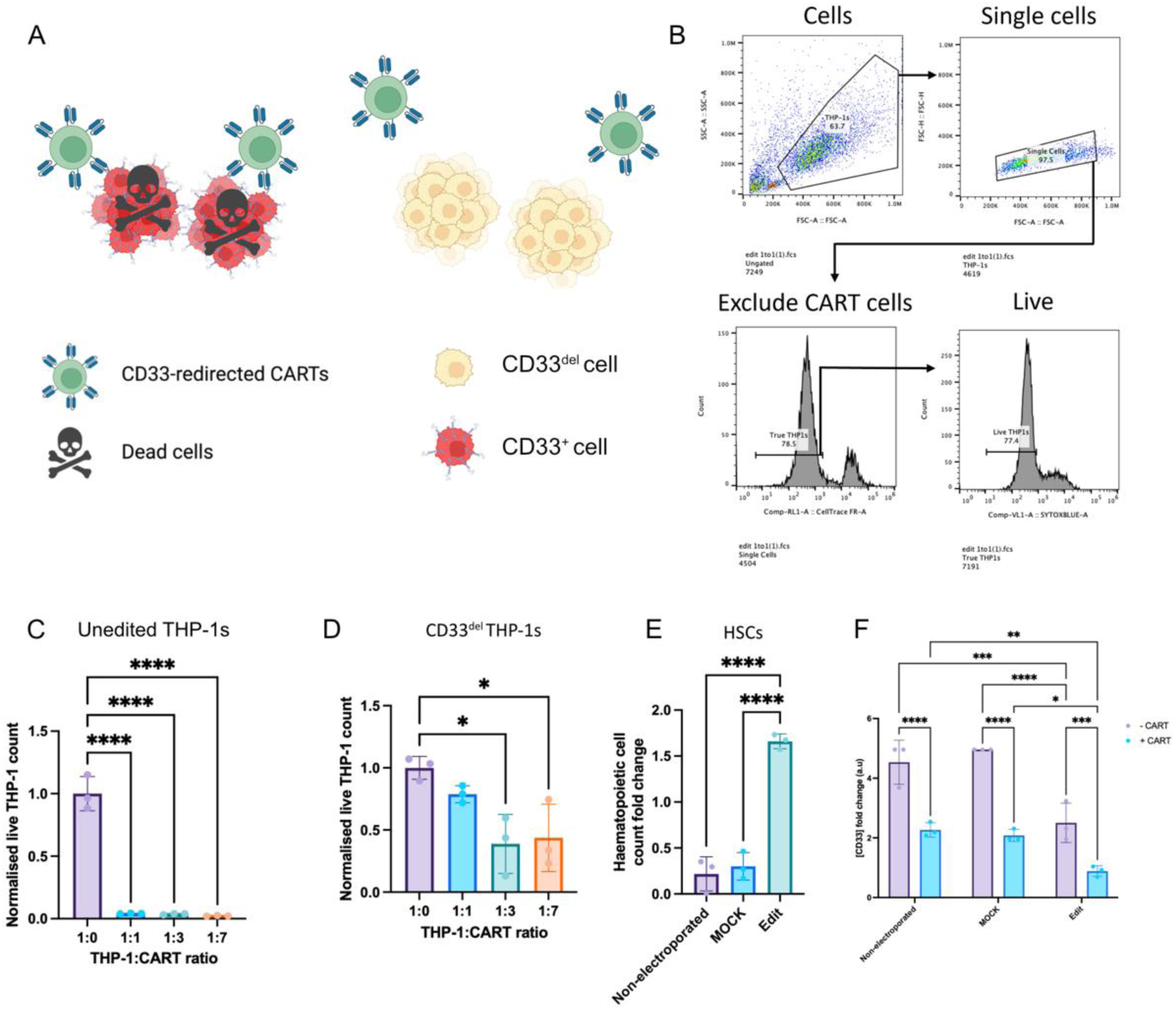
CD33-redirected CAR T-cells selectively ablate CD33+ and not CD33del cells. **A** Illustration of hypothesis that CD33^del^ cells would be spared by CD33-redirected CAR T-cells, while unedited CD33+ cells would be targeted and killed. Created with BioRender.com. **B** Flow cytometry gating strategy for assessing the effect of CAR T-cells in simple two cell type co-culture. Effect of CAR T-cells on **C** unedited and **D** edited THP-1 cell count at various T:E ratios, normalised against no CAR T-cell (1:0) controls, assessed by flow cytometry. **E** Effect of a 1:1 T:E ratio of CAR T-cells on non-electroporated, MOCK and edited HSCs. **F** CD33 expression (MFI) fold change compared to day 0 measurements when CAR T-cells were included with unedited or CD33^del^ HSCs, assessed by flow cytometry. Graphs show mean ± SD. Statistics for **C-E** by one-way ANOVA followed by Tukey multiple comparison test, for **F** by two-way ANOVA followed by Šídák multiple comparison test. *p < 0.05, **p < 0.01, ***p < 0.001 and ****p < 0.0001, non-significant not shown. n=3 experimental replicates for all graphs.

### CRISPR-CAR T-cell therapy in a model BM niche

Next, the CRISPR-CAR T-cell therapy was applied to BM niche models (Fig. 6A). The systems assembled were PEA-FN-BMP2 +MSCs +PeptiGel and liquid cultures with no coatings, MSCs or gel. Combinations of CAR T-cells, THP-1s (used here as an AML model cell line) and unedited or CD33^del^ HSCs were also used. The effect of these cell co-culture combinations on THP-1 cell count was assessed by flow cytometry (Fig 6C). The inclusion of CAR T-cells in all systems was shown to cause a significant decrease in THP-1 count across conditions, demonstrating successful on-target killing of CD33+ AML cells in complex niche systems. Also, the addition of HSCs in the niche caused an unexpected downwards trend in THP-1 count.

**Figure 6.**
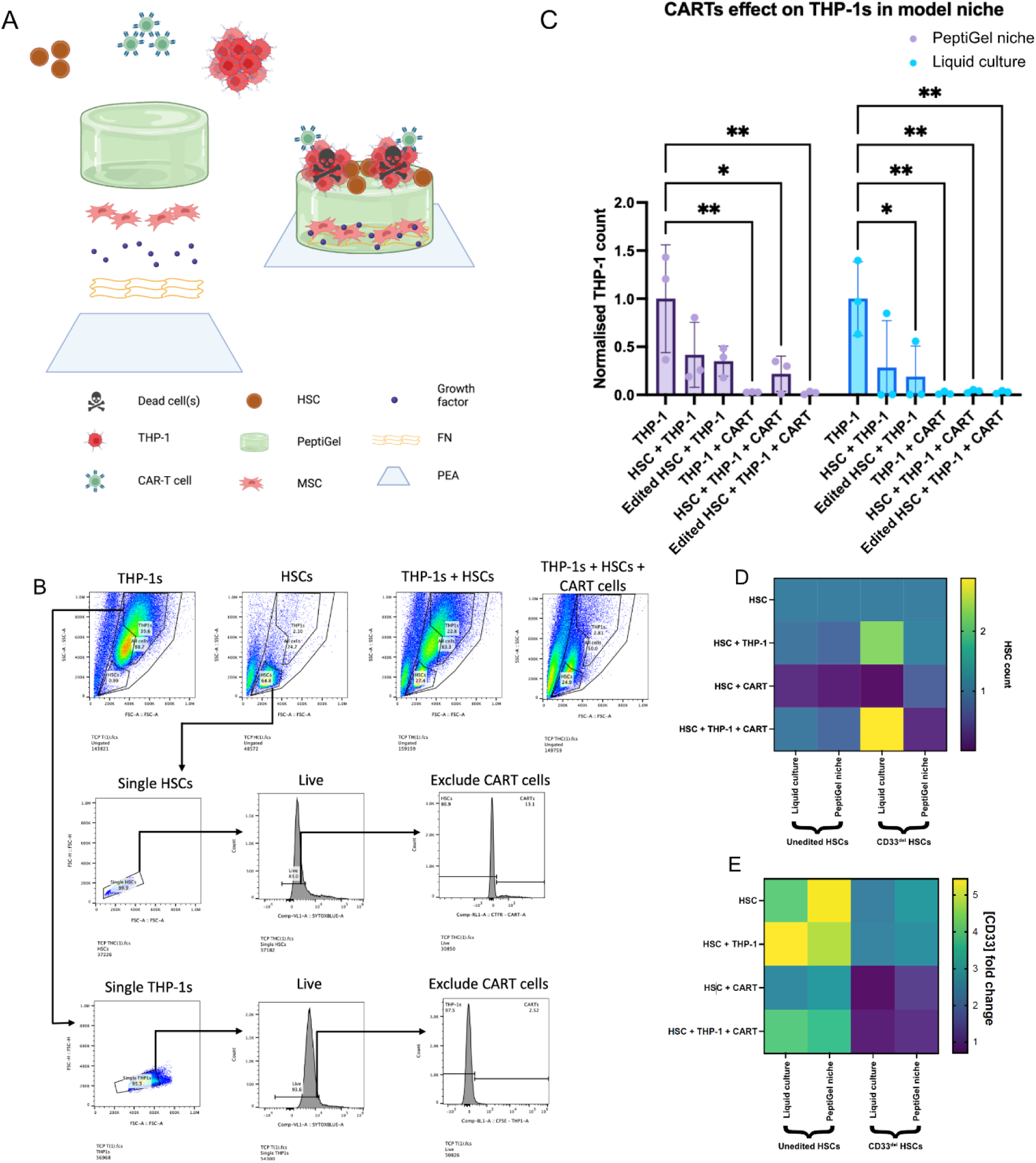
Effect of CD33-redirected CAR T-cells on CD33^del^ cells in niche systems. **A** Schematic of the synthetic BM model for testing CAR T-cells. Created with BioRender.com. **B** Flow cytometry gating strategies for assessing the effect of CAR T-cells in complex co-cultures, with representative plots showing how different cell types were identified with SSC-A and FSC-A, and how the inclusion of CAR T-cells affected the cell populations. **C** Effect of CAR T-cells on THP-1 cell count in HSC/CD33^del^ HSC/no co-culture conditions assessed by flow cytometry. The PeptiGel niche included pPEA, FN, BMP2, MSCs, PeptiGel and the cell types listed for each condition. The liquid culture control conditions were uncoated PCS plates without MSCs and with the cell types listed. **D** Effect of different cell combinations, niche systems and editing status of HSCs on HSC count normalised against HSC only controls for each condition assessed by flow cytometry. **E** Effect of different cell combinations, niche systems and editing status of HSCs on HSC CD33 expression (MFI) assessed by flow cytometry. Graph shows mean ± SD, heat maps show mean. Statistics for **C** by two-way ANOVA followed by Tukey multiple comparisons test. *p < 0.05, **p < 0.01, ***p < 0.001 and ****p < 0.0001 non-significant not shown. **C-E** n=3 experimental replicates.

The effect of CAR T-cells on unedited HSC and CD33^del^ HSC counts in niche models was also assessed (Fig. 6D). The HSC count was stable when THP-1s were included and expectedly decreased in response to the addition of CAR T-cells in all conditions, except when CD33^del^ HSCs were cultured in a PeptiGel system. The level of CD33 expression of HSCs in each system was also assessed to further investigate the observed effect and whether it was CD33-dependant (Fig. 6E). The inclusion of CAR T-cells appeared to cause a drop in CD33 expression, which was additive with the effect of the CRISPR-induced CD33 knock-out.

## Discussion

In this study we aimed to develop a biomimetic BM niche model for testing novel therapies for BM-associated diseases and disorders. Current platforms rely on the use of animal-derived components that can introduce variability into systems. Therefore we sought to mimic the elastic nature of the BM ECM using a fully synthetic hydrogel platform to control an HSC-supportive phenotype in MSCs. We then introduced AML cells into the platform and investigated the on-target efficacy of a combined CRISPR-CAR T-cell therapy for AML.

We first employed the PEA-FN-BMP2 nanoscale coating system on PCS tissue culture plates to support a physiological ECM-like surface (Fig. 1). The PEA system caused FN to adopt an open conformation, allowing MSCs to interacted with the exposed FN RGD domains with integrins [64], and the BMP2 bound to the GF binding domain with BMP receptor complexes [65,66] in a synergistic manner [67]. The mechanical properties of fully synthetic, peptide-based, viscoelastic PeptiGels were also assessed and found to be similar to BM [46,47,68].

Then, a system combining the ECM-like surface and PeptiGels was used to support a nestin+ SCF+ MSC niche-like phenotype that could act as an HSC-supportive feeder layer (Fig. 2). The addition of the PeptiGel also acted as a barrier between the MSCs and HSCs, facilitating analysis of these cell populations independently [26]. The PeptiGels were stiffer (∼1000 Pa with no media, ∼4500 Pa with 2% DMEM) than previously utilised collagen gels (100 Pa), thus in combination with the stiff polymer surface the cells were cultured on, may have replicated the stiffer bone-lining region of the BM niche rather than the soft central region of the niche potentially replicated by the collagen gel systems [48,49].

Next, we assessed the ability of our optimised niche to maintain a population of HSCs in vitro (Fig. 3). We found that the inclusion of MSCs significantly improved haematopoietic cell, HSC and LT-HSC numbers across conditions, similarly to the endogenous niche [69,70], highlighting the usefulness of an MSC feeder layer for in vitro HSC culture [26]. In addition, the increased fold change in LT-HSC numbers when MSCs were included in Peptigel systems compared to collagen and no gel controls suggested that the MSCs in the PeptiGel system acted as a superior feeder layer. However, the total number of haematopoietic cells was much lower in PeptiGel systems than the collagen and no gel systems, even with the inclusion of MSCs. This was likely due to other systems driving proliferation of the haematopoietic compartment, whereas the lack of biological milileu in the Peptigel systems may have led to quiescence [71]. The differentiation and phenotype of haematopoietic cells was also affected by the niche model they were cultured in. Increased megakaryopoiesis was observed in the PeptiGel niches. Recent research has suggested that increased megakaryocyte differentiation is the native state of unperturbed haematopoiesis, and that the classical model seen in most in vitro and animal models is a stress-induced, atypical pathway [72]. Therefore, the PeptiGel model could have been encouraging haematopoietic differentiation to occur as it does when undisturbed in vivo. As a result of these findings, we concluded that our niche was able to sufficiently support HSC cell maintenance in vitro by replicating aspects of the BM niche microenvironment.

We then aimed to develop a combined CRISPR-CAR T-cell system to target AML. This approach, in which CD33 is deleted from healthy HSCs that are then transplanted alongside CD33-redirected CAR T-cells to allow sustained myelopoiesis despite long-term ablation of CD33^+^ cells, has been successfully demonstrated in simple in vitro systems and animal models [62,73]. However, while there are examples of CAR T-cell therapy and curative HSC gene editing therapies advancing to clinical trial [74,75], to our knowledge a combined approach has yet to reach this stage. Therefore, this novel therapeutic approach is perfectly positioned to investigate with our system. To effectively disrupt CD33 in HSCs we developed a cocktail of three sgRNAs targeting regions of DNA within or associated with the CD33 gene [62] (Fig. 4). We demonstrated at a protein and DNA level that this approach increased the efficiency of the gene edit compared to sgRNAs alone. In HSCs, this was due to a lack of CD33 accumulation on edited CD33^del^ cells compared to unedited control cells. This was likely due to differentiation of both cell populations throughout the experiment, demonstrated by increased CD13 and lineage marker expression, resulting in accumulation of CD33 on unedited cells due to myeloid differentiation that was prevented by the edit on CD33^del^ cells. Of note, CD33 expression appears to rise in CD33^del^ HSCs before falling again, which could be attributed to cytosolic CD33 precursors present prior to editing maturing and translocating to the cell membrane but being diluted with subsequent mitotic events and a lack of replenishment due to the edit. The improved efficiency of the sgRNA cocktail compared to single sgRNA-induced edits may have been due to the introduction of double strand breaks caused by the CRISPR-Cas9 system at multiple sites within single cells, possibly overwhelming the DNA repair mechanisms. Using this approach, we were able to produce highly efficient disruption of CD33 in HSCs.

Next, we investigated how the gene edit we had created affected the sensitivity of cells to CD33-redirected CAR T-cells. Initially we tested this system in simple liquid co-culture (Fig. 5) and found that the edit gave cells significant protection from CAR T-cell-induced death, completely preventing a drop in CD33^del^ HSC count when CAR T-cells were present, despite a clear drop being present for unedited cells. Interestingly, we also observed a drop in CD33 expression across conditions when CAR T-cells were included. In unedited controls this was assumed to be due to ablation of CD33^+^ cells, resulting in a surviving population of cells with reduced CD33 expression. However, this doesn’t explain the drop in CD33 expression seen in CD33^del^ HSCs co-cultured with CAR T-cells, as no corresponding drop in cell count was observed. The drop in CD33 may therefore have been due to antigen loss by HSCs expressing low levels of CD33 due to incomplete knock-out/knock-down in response to the presence of CAR T-cells, which has been reported previously in other cell types [63,76]. This phenomenon could result in differences in HSC differentiation [76,77], which may be a confounding issue for this therapeutic approach if it prevented the reconstruction of a normal haematopoietic system by CD33^del^ HSCs in the presence of CAR T-cells. Our results therefore shed light on some of the potential issues with this therapeutic approach and highlight the effectiveness of our gene edit for protecting HSCs from CD33-redirected CAR T-cells.

Finally, we tested the efficacy of the CRISPR-CAR T-cell therapy in the model BM niche (Fig. 6). We showed that even in this highly complex environment, AML cells were still effectively eliminated by CAR T-cells, demonstrating the promise that this therapeutic approach holds. In addition, a drop in AML cell count was observed when co-cultured with HSCs across conditions. This suggested that we modelled the graft vs leukaemia GvL effect. This effect, which is essential to the function of curative HSC transplants for leukaemia, is caused by transplanted HSCs displacing endogenous AML cells, which they recognise as non-self, by producing immune cells to target them and competing for resources [78]. This effect has not to our knowledge been modelled in vitro previously, demonstrating the possibility of modelling highly complex interactions in vitro using systems as presented here. Notably, CAR T-cell addition in model BM niches and simple 2D co-culture systems caused a drop in HSC numbers, even when the HSCs were edited CD33^del^ cells. This may have been due the short time span of this experiment, which could have resulted in higher levels of surface CD33 protein in HSCs due to maturing cytosolic CD33, as discussed previously. Despite this, across conditions, the level of CD33 expression dropped when HSCs were edited and when they were co-cultured with CAR T-cells, and this response was additive. This implied that the edit was working and that CD33^+^ cells were being preferentially eliminated, possibly supplemented by antigen loss. Therefore, if this system can be further optimised and the timings of CD33 expression and maximal engraftment capacity investigated, it holds significant promise. The niche model presented here, and other bioengineered systems could provide the ideal platform for this optimisation. Furthermore, when THP-1s were included alongside HSCs and CAR T-cells, they appeared to have a protective effect across conditions. This may have been due to the CAR T-cells preferentially targeting the THP-1 AML cells, which express high levels of CD33, and becoming exhausted in the process [79]. This shielding could be used to further reduce off-target effects of CD33-redirected CAR T-cells if selectively applied to patients with AML cells that strongly express CD33. The usefulness of the model BM niche for dissecting novel therapeutic approaches was also demonstrated by its role in this research.

In conclusion, we have successfully developed a simple, synthetic BM niche which replicates some of the complexity of the endogenous niche. Our niche facilitated testing of a CRISPR-CAR T-cell therapy for AML which has been researched extensively but has yet to reach clinical trial. This therapy worked well in standard tissue culture conditions. When tested in the synthetic niche however, this therapeutic approach was shown to be highly complex, causing ablation of HSCs that received a protective gene edit, as well as AML cells. These results highlight the usefulness of *in vitro* organ models as a cruelty-free, fully human alternative for bridging the gap between pre-clinical research and clinical trials.

## Materials and methods

### Preparation of materials

96 well plates with polymer coverslip bottoms (Ibidi, 89626) or standard tissue culture plastic plates were placed in a bespoke plasma chamber perpendicular to the plasma flow. Samples were exposed to air plasma for 5 minutes at 50W of radio frequency (RF) incident power to ensure removal of any residual organic matter. Samples were then coated with PEA (Sigma-Aldrich, E9706) using an RF power of 50W. The plasma treatment was carried out for 15 minutes[31,39,80,81]. Plates were then sterilised by irradiating them with UV light for 30 minutes. FN (Sigma-Aldrich, F2006-2MG) diluted to 20 µg/mL in sterile phosphate buffered saline (PBS) (Gibco, 14190-094) was added to wells. Plates were incubated at room temperature (RT) for 1h before the FN solution was removed and the wells were washed twice with PBS. BMP2 (Sigma-Aldrich, H4791-10UG) diluted to 50 ng/mL in sterile PBS was added to wells and incubated for 1h at RT. Plates were then washed twice with PBS.

### MSC culture

MSCs (PromoCell, C-12974) were incubated in a humidified 5% CO_2_ atmosphere at 37°C. When expanding MSCs, they were cultured in Dulbecco’s Modified Eagle’s Medium (DMEM, Sigma-Aldrich) with 10% v/v FBS (Thermo Fisher), 1% v/v nonessential amino acids (Gibco, 11140-035), 1% v/v 100 mM sodium pyruvate (Sigma-Aldrich, S8636-100ML) and 3% v/v of an antibiotic mix consisting of 10 mg/ml penicillin/streptavidin (Gibco, 15140-122), 200 nM L-glutamine (Gibco, 25030-024), and 0.5% v/v amphotericin (Gibco, 15290-026), later described as 10% DMEM. When using plates, MSCs were seeded at 3×10^3^ cells/cm^2^ and cultured using identical media containing but with 2% FBS, later described as 2% DMEM.

### HSC culture

HSCs were isolated from the bone marrow aspirates of patients undergoing joint replacement surgery. 30 patient samples were used with patients ranging from 26 to 91 years old, with a mean age of 62. 55% of samples were from male patients, 45% female. Mononuclear cells were separated by density gradient centrifugation using Ficoll-Paque. Following nucleated cell isolation, CD34+ cells were isolated using a EasySep^TM^ Human CD34 Positive Selection Kit II (Stemcell technologies) according to the manufacturer’s protocol. These cells were then counted using a haemocytometer before reseeding 5×10^4^ cells/mL in IMDM+ in a 24 well plate and incubating overnight. Alternatively, the cells were frozen immediately after EasySep separation using HSC freezing media. For the experiments used to produce figures 3, 5 and 6, CD34+ HSCs were purchased (Stemcell technologies, 70002), and one sample used per experiment per biological repeat.

### THP-1 culture

Non-adherent THP-1 cells (DSMZ, ACC16) were seeded in flasks at ∼ 5×10^5^ cells/mL. 2-3 times per week when the cells reached confluency ∼80% of cell suspension was removed and replaced with fresh 10% RPMI.

### Gel assembly

Collagen type I gel was made on ice by combining 2.5 mL type-1 rat tail collagen (First Link, 60-30-810), 0.5 mL FBS (Sigma-Aldrich, F9665-500ML), 0.5 mL 10X DMEM (Sigma, D2429-100ML) and 0.5 mL 10% DMEM, adjusted to pH 8.2 using sodium hydroxide. Collagen gel was then added to plates and polymerised overnight at 37°C 5% CO_2_. Note that PeptiGel was delivered as pre-made gels.

### Rheology

A Discovery Hybrid 2 (DHR-2) rheometer (TA instruments) was used to measure the mechanical properties of PeptiGel and collagen. A 20 mm diameter parallel plate geometry rheometer head was used, with the geometry gap set to 500 μm and the trim gap set to 50 μm. All measurements were proceeded by a 3 min temperature soak at the desired temperature. Amplitude sweeps were carried out at 1.0 Hz, 0.05-40% strain. To test stress recovery, the gels were subjected to 0.2% strain at 1.0 Hz for 5 minutes. They were then placed under 100% strain at 1.0 Hz for a further 5 minutes. This process was repeated four times in total. Young’s modulus was calculated using the G^’^ from the linear region and multiplying by 3, assuming a Poisson’s ratio of 0.5 in accordance with Hooke’s law. Tan delta was calculated as G’ ÷ G”.

### Assembling model BM niche

Following plate preparation, MSCs (PromoCell, C-12974) were seeded in 2% DMEM in plates and cultured overnight at 37°C. The following day all media was removed from plates. 70 μL of PeptiGel (Cell Guidance Systems) was added using a positive displacement pipette before the plate was centrifuged at 333 g for 1 min. For collagen gel controls, 70 μL of collagen was then added to appropriate wells. 2% DMEM was added to the wells not containing collagen. Within 30 min of incubation starting, half of the media in wells without collagen was replaced with fresh 2% DMEM twice. Plates were incubated overnight at 37°C in 5% CO_2_. After 1 day half of the media on each well was replaced with fresh media, and media was added to collagen wells. Another half media change was performed on day 3. If the system was to be used for co-culture, 7 days after cell seeding, all media was removed and 7.5×10^3^ of each cell type used for co-culture were added in appropriate media (for THP-1s, Roswell Park Memorial Institute (RPMI) (Gibco, 21875-034) with 10% v/v FBS and 2% v/v of the previously described antibiotic mix, later described as 10% RPMI. For all other co-cultures, Iscove Modified Dulbecco Media (IMDM) (Merck, I3390-500ML) with 1.96 mM L-glutamine (Gibco, 25030-024), 1% v/v penicillin/streptomycin, 20% bovine insulin transferrin (BIT) (Stemcell technologies, 09500), 0.1% fms-like tyrosine kinase 3 ligand (FL) (Peprotech, 300-19), 0.1% SCF (Peprotech, 300-07) and 0.05% thrombopoietin (TPO) (Peprotech, GMP300-18), later described as IMDM+). If assessing non-adherent cells with flow cytometry, cells were harvested 5 days after addition to the niche and analysed immediately.

### XPS analysis

Spectra were obtained at the National EPSRC XPS Users’ Service at Newcastle University. XPS was performed using a K-Alpha apparatus (Thermo Scientific), with a microfocused monochromatic Al Kα source (X-ray energy = 1486.6 eV) at a voltage of 12 kV, current of 3 mA, power of 36 W, and spot size of 400 µm × 800 µm. Spectra analysis and curve fitting were performed using CasaXPS software.

### AFM

Plates were coated with PEA and/or FN or left uncoated, then individual wells were cut out using a scalpel and/or wire cutters. If stored, surfaces were covered with PBS and kept at 4°C for up to a week. Prior to imaging, surfaces were dried with high pressured nitrogen, then imaged with a Drive AFM apparatus from Nanosurf in Dynamic Force mode using a conical cantilever holder, ceramic, grooveless (BT08891), and a 7.3 N/m pyramidal silicon cantilever from apex probes (USC-F1.2-K7.3-10); amplitudes were driven at the resonance frequency of the cantilever (∼1200 kHz). Height images were acquired from each scan using the following parameters: scan frequency of 1 Hz, 500 points per line, 1 µm^2^ scan area, 30% amplitude reduction from the maximum free vibration amplitude. Gwyddion software was used to prepare and analyse AFM images. Image data was levelled by mean plane subtraction, then to make facets point upwards. Rows were then aligned using line medians, and horizontal scars were corrected. The processed image was then used to determine mean roughness (Sa).

### FN in cell western

Surfaces were coated as described, blocked with 1% w/v bovine serum albumin (BSA) (Sigma-Aldrich, A7906-100G). Primary antibodies were added in 1% w/v BSA overnight at 4°C: a mouse polyclonal anti-FN antibody (Sigma-Aldrich, F3648) diluted 1:400, a mouse monoclonal anti-P5F3 antibody (Santa Cruz, sc-18827) diluted 1:100, or a mouse monoclonal anti-HFN7 antibody (abcam, ab212371) diluted 1:100. Wells were washed 3×5 min with 0.5% v/v Tween20 (PBST). Secondary antibodies were added in 1% w/v BSA for 1 h at 37°C: goat anti-rabbit channel 800 antibody (Li-Cor, 926-32211) at a 1:800 dilution, and a goat anti-mouse channel 680 antibody (Li-Cor, 926-68070) at a 1:800 dilution. Plates were washed 3×5 min again with PBST, then imaged using a Li-Cor Odyssey M to quantify immunofluorescence.

### Immunofluorescence microscopy

If cells were being imaged for excreted proteins such as SCF or CXCL12, ∼24 h before fixation the media in wells to be analysed was replaced with identical media containing 5 µg/mL of brefeldin A (SigmaAldrich, B6542). MSCs were processed 7 days after being added to wells. Cells underwent fixation with 10% v/v formaldehyde (Fisher scientific, F/1501/PB17), 2% w/v sucrose (Fisher scientific, S/8600/60) for 30 min at 37°C. If PeptiGel was included, it was removed with two washes of ice-cold PBS. Cells were permeabilised for 10 min at RT with 0.5% v/v Triton-X (Sigma-Aldrich, T9284-100ML), 301 mM sucrose (Fisher scientific, S/8600/60), 5 mM sodium chloride (VWR chemicals, 27810.295), 2.95 mM magnesium chloride hexahydrate (VWR chemicals, 25108.260), 20 mM hepes (VWR chemicals, 441485H). Cells were blocked for 1 h at 37°C using 1% w/v BSA). Primary antibodies were added in 1% w/v BSA and stored on a shaker at 4°C overnight. The antibodies were: Nestin (Proteintech 19483-1-AP) 1:100, SCF (abcam ab64677) 1:200, SDF1 (aka CXCL12) (abcam ab155090) 1:100. After 3×5 min washes at RT with PBST, secondary antibodies in 1% w/v BSA were added for 1h at 37°C: horse anti-mouse biotinylated secondary antibody (Vector laboratories, BA-2000) 1:50. Phalloidin (Invitrogen, O7466) diluted 1:200 was also included in the primary antibody solution to stain actin. 3×5 min washes were performed with PBST at RT, and rhodamine streptavidin (Invitrogen, R415) in 1% w/v BSA was added 1:50 at 4°C for 30 min, before plates were washed 3×5 min again. 1 drop per mL of NucBlue live cell stain (Invitrogen, R37605) in PBS was added for 30 min at RT before 3×5 min final washes. Plates were stored at 4°C in foil for up to two weeks prior to imaging. Protein expression was assessed using a custom FIJI macro for corrected total cell fluorescence (CTCF) analysis.

### Flow cytometry

Only non-adherent cells were analysed by flow cytometry. Cells were harvested by extracting cell suspension, washing wells with fresh media, and extracting washes. Cells were centrifuged at 387g for 5 min, then stained on ice for 45 min in flow buffer (0.5% w/v BSA, 2 mM Ethylenediaminetetraacetic acid (EDTA) (Invitrogen, 15575-038) in PBS) containing appropriate antibodies. For basic HSC characterisation: sytox blue (ThermoFisher scientific, S34857), CD45 (Miltenyi Biotec, 130-110-635), Lineage cocktail (Lin) (ThermoFisher scientific, 22-7778-72), CD34 (Miltenyi Biotec, 130-113-176). For LT-HSC characterisation: sytox blue (ThermoFisher scientific, S34857), CD45 (Miltenyi Biotec, 130-110-635), Lin (ThermoFisher scientific, 22-7778-72), CD34 (Miltenyi Biotec, 130-113-176), CD38 (ThermoFisher scientific, 25-0388-42), CD90 (Invitrogen, 56-0909-42), CD45RA (ThermoFisher scientific, 63-0458-42). For extended haematopoietic cell characterisation: sytox blue (ThermoFisher scientific, S34857), CD45 (Miltenyi Biotec, 130-110-635), CD16 (ThermoFisher scientific, 69-0168-42), CD41/61 (ThermoFisher scientific, MA5-44123), CD7 (ThermoFisher scientific, 46-0078-42), CD34 (Miltenyi Biotec, 130-113-176), CD38 (ThermoFisher scientific, 25-0388-42), CD90 (Invitrogen, 56-0909-42), CD45RA (ThermoFisher scientific, 63-0458-42). For THP-1s: sytox blue (ThermoFisher scientific, S34857), CD34 (Miltenyi Biotec, 130-113-176), CD38 (ThermoFisher scientific, 25-0388-42). For determining CD33 expression: sytox blue (ThermoFisher Scientific, S34857), CD33 HIM3-4 domain (Invitrogen, 15-0339-42), CD33 WM53 domain (BioLegend, 303421). When CAR T-cells were included, CAR T-cells were stained with cell trace far red (CTFR) (Invitrogen, C34572) before co-culture. Other stains included for CAR T-cell effect analysis: sytox blue (ThermoFisher Scientific, S34857), CD33 HIM3-4 domain (Invitrogen, 15-0339-42), CD33 WM53 domain (BioLegend, 303421). The cells were washed with flow buffer then analysed with an attune NxT acoustic focusing cytometer. Gating strategies are shown in relevant figures.

### CRISPR

Ribonucleoprotein (RNP) complexes were prepared: 1.5 µM Cas9 (IDT, 1081060), 3.85 µM electroporation buffer (IDT, 1075916), 2% v/v glycerol (Fisher scientific, G/0650/17), 2.25 µM sgRNA (CD33 exon 2 – GGGGAGUUCUUGUCGUAGUA, CD33 exon 3 – CCUGUGGGUCAAGUCUAGUG (from [62]), CD33 upstream enhancer – GAUACAAGCAGACCACCAGA) in RNAse-free water. If multiple sgRNA were included, the total concentration of sgRNA was divided equally between sgRNAs. ∼2.4×10^5^ cells were electroporated (THP-1s: 1600 V/10 ms/3 pulses, HSCs: 1700 V/20 ms/1 pulse) using a Neon transfection system (ThermoFisher scientific).

### TIDE analysis

Cells’ gDNA was isolated using a DNeasy blood & tissue kit (Qiagen). sgRNA target sites were PCR amplified using a PCR master mix: 1Xphusion HF buffer, 200 µM dNTP mix, 3% v/v dimethylsulfoxide, 20 U/mL Phusion HF polymerase, nuclease free water (all NEB, EO553S), 500 nM primers (CD33 exon 2 F primer – AGCTGCTTCCTCAGACATGC (from [73]), CD33 exon 2 R primer – CAGGGATGAGGATTTTGGGC, CD33 exon 3 F primer – GGGAAGTTCATGGGTACTGC, CD33 exon 3 R primer – CATCCTGTCTCCCCTACACC, CD33 enhancer F primer – TGAAAGGCATGCACTCAGAA, CD33 enhancer R primer – TATCCAGCCCCAAATGCCA). PCR products were processed using a QIAquick PCR Purification Kit (Qiagen) and sanger sequenced (Eurofins Genomics). Mutagenesis analysis was done using TIDE analysis [82] (http://shinyapps.datacurators.nl/tide/).

### CAR T-cell culture and staining

CD33-redirected CAR T-cells (CD33 scFv-Beam 2-TM-CD28-CD3z) (AMSbio, AMS.PM-CAR1056-1M) were cultured in 10% RPMI. When staining with CTFR, CAR T-cells were pelleted then resuspended in 1mL 10% RPMI with 1 nM CTFR (Invitrogen, C34572) for 20 minutes at 37°C. 15mL 10% RPMI was then added, and the suspension incubated for a further 5 minutes at 37°C. Afterwards, the suspension was centrifuged at 300 x g, then resuspended in 1 mL of media used in the experiment the CAR T-cells were being added to and incubated for 10 minutes at 37°C.

## Data availability

The raw and processed data presented will be made available upon final publication.

## Acknowledgements

The author would like to thank Jennifer Cassels at the Paul O’Gorman Leukaemia Research Centre for conducting fluorescence assisted cell sorting, as well as Connor Robinson, Emma Kelly, Yinbo Xiao and Lauren Hope for providing cells.

## Author contributions

W. Sebastian Doherty-Boyd: conceptualisation, data curation, formal analysis, investigation, methodology, project administration, software, validation, visualisation, writing – original draft presentation, writing – review and editing. P. Monica Tsimbouri: conceptualisation, methodology, supervision. Vineetha Jayawarna: formal analysis, investigation, validation. Matthew Walker: investigation, validation. Aqeel F. Taqi: methodology. Niall Mahon: methodology. Dominic Meek: resources. Peter Young: resources. Aline Miller: resources, supervision. Adam West: methodology, resources, supervision. Manuel Salmeron-Sanchez: resources, supervision. Matthew J Dalby: conceptualisation, methodology, project administration, resources, supervision. Hannah Donnelly: conceptualisation, methodology, supervision, writing – review and editing.

## References

[1] Vetrie D, Helgason GV, Copland M. The leukaemia stem cell: similarities, differences and clinical prospects in CML and AML. Nat Rev Cancer 2020;20:158–73. 10.1038/s41568-019-0230-9.

[2] Pinho S, Frenette PS. Haematopoietic stem cell activity and interactions with the niche. Nat Rev Mol Cell Biol 2019. 10.1038/s41580-019-0103-9.

[3] Méndez-Ferrer S, Bonnet D, Steensma DP, Hasserjian RP, Ghobrial IM, Gribben JG, et al. Bone marrow niches in haematological malignancies. Nat Rev Cancer 2020;20:285–98. 10.1038/s41568-020-0245-2.

[4] Fry TJ, Mackall CL. T-cell adoptive immunotherapy for acute lymphoblastic leukemia. Hematology / the Education Program of the American Society of Hematology American Society of Hematology Education Program 2013;2013:348–53. 10.1182/asheducation-2013.1.348.

[5] Qin H, Cho M, Haso W, Zhang L, Tasian SK, Oo HZ, et al. Eradication of B-ALL using chimeric antigen receptor-expressing T cells targeting the TSLPR oncoprotein. Blood 2015;126:629–39. 10.1182/blood-2014-11-612903.

[6] Tambaro FP, Singh H, Jones E, Rytting M, Mahadeo KM, Thompson P, et al. Autologous CD33-CAR-T cells for treatment of relapsed/refractory acute myelogenous leukemia. Leukemia 2021;35:3282–6. 10.1038/s41375-021-01232-2.

[7] Park JH, Rivière I, Gonen M, Wang X, Sénéchal B, Curran KJ, et al. Long-Term Follow-up of CD19 CAR Therapy in Acute Lymphoblastic Leukemia. New England Journal of Medicine 2018;378:449–59. 10.1056/nejmoa1709919.

[8] Denlinger N, Bond D. CAR T-cell therapy for B-cell lymphoma 2022;46:1–16. 10.1016/j.currproblcancer.2021.100826.CAR.

[9] Atilla E, Benabdellah K. The Black Hole: CAR T Cell Therapy in AML. Cancers (Basel) 2023;15:1–19. 10.3390/cancers15102713.

[10] Robinson S, Finel H, Boumendil A, Tilly H, Salles G, Corradini P, et al. Remissions of Acute Myeloid Leukemia and Blastic Plasmacytoid Dendritic Cell Neoplasm Following Treatment with CD123-Specific CAR T Cells: A First-in-Human Clinical Trial. Blood 2017;130:811. 10.1182/blood.V130.Suppl.

[11] Zhang H, Bu C, Peng Z, Li G, Zhou Z, Ding W, et al. Characteristics of anti-CLL1 based CAR-T therapy for children with relapsed or refractory acute myeloid leukemia: the multi-center efficacy and safety interim analysis. Leukemia 2022;36:2596–604. 10.1038/s41375-022-01703-0.

[12] Marofi F, Rahman HS, Al-Obaidi ZMJ, Jalil AT, Abdelbasset WK, Suksatan W, et al. Novel CAR T therapy is a ray of hope in the treatment of seriously ill AML patients. Stem Cell Res Ther 2021;12:1–23. 10.1186/s13287-021-02420-8.

[13] Kim MY. Making normal hematopoiesis invisible to CAR T cells. Trends Cancer 2023;9:983–4. 10.1016/j.trecan.2023.10.001.

[14] Molica M, Perrone S, Mazzone C, Niscola P, Cesini L, Abruzzese E, et al. Cd33 expression and gentuzumab ozogamicin in acute myeloid leukemia: Two sides of the same coin. Cancers (Basel) 2021;13:1–20. 10.3390/cancers13133214.

[15] Van Der Velden VHJ, Te Marvelde JG, Hoogeveen PG, Bernstein ID, Houtsmuller AB, Berger MS, et al. Targeting of the CD33-calicheamicin immunoconjugate Mylotarg (CMA-676) in acute myeloid leukemia: In vivo and in vitro saturation and internalization by leukemic and normal myeloid cells. Blood 2001;97:3197–204. 10.1182/blood.V97.10.3197.

[16] Laszlo GS, Estey EH, Walter RB. The past and future of CD33 as therapeutic target in acute myeloid leukemia. Blood Rev 2014;28:143–53. 10.1016/j.blre.2014.04.001.

[17] Gill S, Tasian SK, Ruella M, Shestova O, Li Y, Porter DL, et al. Preclinical targeting of human acute myeloid leukemia and myeloablation using chimeric antigen receptor– modified T cells. Blood 2014;123:2343–54. 10.1182/blood-2013-09-529537.

[18] Kim MY, Yu KR, Kenderian SS, Ruella M, Chen S, Shin TH, et al. Genetic Inactivation of CD33 in Hematopoietic Stem Cells to Enable CAR T Cell Immunotherapy for Acute Myeloid Leukemia. Cell 2018;173:1439–1453.e19. 10.1016/j.cell.2018.05.013.

[19] Siegler EL, Wang P. Preclinical Models in Chimeric Antigen Receptor-Engineered T-Cell Therapy. Hum Gene Ther 2018;29:534–46. 10.1089/hum.2017.243.

[20] Kiesgen S, Messinger JC, Chintala NK, Tano Z, Adusumilli PS. Comparative analysis of assays to measure CAR T-cell-mediated cytotoxicity. Nat Protoc 2021;16:1331–42. 10.1038/s41596-020-00467-0.

[21] Duncan BB, Dunbar CE, Ishii K. Applying a clinical lens to animal models of CAR-T cell therapies. Mol Ther Methods Clin Dev 2022;27:17–31. 10.1016/j.omtm.2022.08.008.

[22] Ingber DE. Human organs-on-chips for disease modelling, drug development and personalized medicine. Nat Rev Genet 2022;23:467–91. 10.1038/s41576-022-00466-9.

[23] Abolins S, King EC, Lazarou L, Weldon L, Hughes L, Drescher P, et al. The comparative immunology of wild and laboratory mice, Mus musculus domesticus. Nat Commun 2017;8. 10.1038/ncomms14811.

[24] Michaels YS, Buchanan CF, Gjorevski N, Moisan A. Bioengineering translational models of lymphoid tissues. Nature Reviews Bioengineering 2023;1:731–48. 10.1038/s44222-023-00101-0.

[25] Ingber DE. Is it Time for Reviewer 3 to Request Human Organ Chip Experiments Instead of Animal Validation Studies? Advanced Science 2020;2002030:1–15. 10.1002/advs.202002030.

[26] Doherty-Boyd WS, Donnelly H, Tsimbouri MP, Dalby MJ. Building bones for blood and beyond: the growing field of bone marrow niche model development. Exp Hematol 2024;135:104232. 10.1016/j.exphem.2024.104232.

[27] Rico P, Mnatsakanyan H, Dalby MJ, Salmerón-sánchez M. Material-Driven Fibronectin Assembly Promotes Maintenance of Mesenchymal Stem Cell Phenotypes 2016:6563–73. 10.1002/adfm.201602333.

[28] Donnelly H, Kurjan A, Yong LY, Xiao Y, Lemgruber L, West C, et al. Fibronectin matrix assembly and TGFβ1 presentation for chondrogenesis of patient derived pericytes for microtia repair. Biomaterials Advances 2023;148. 10.1016/j.bioadv.2023.213370.

[29] Xiao Y, Donnelly H, Sprott M, Luo J, Jayawarna V, Lembruger L, et al. Material-driven fibronectin and vitronectin assembly enhances BMP-2 presentation and osteogenesis. Mater Today Bio 2022:100310. 10.1016/j.mtbio.2022.100367.

[30] Llopis-hernández V, Cantini M, González-garcía C, Cheng ZA, Yang J, Tsimbouri PM, et al. Material-driven fibronectin assembly for high-efficiency presentation of growth factors. Sci Adv 2016:1–11. 10.1126/sciadv.1600188.

[31] Cheng ZA, Alba-Perez A, Gonzalez-Garcia C, Donnelly H, Llopis-Hernandez V, Jayawarna V, et al. Nanoscale Coatings for Ultralow Dose BMP-2-Driven Regeneration of Critical-Sized Bone Defects. Advanced Science 2018;1800361:1800361. 10.1002/advs.201800361.

[32] Bowerman CJ, Nilsson BL. Self-assembly of amphipathic β-sheet peptides: insights and applications. Biopolymers 2012;98:169–84. 10.1002/bip.22058.

[33] Mohammed A, Miller AF, Saiani A. 3D networks from self-assembling ionic-complementary octa-peptides. Macromol Symp 2007;251:88–95. 10.1002/masy.200750512.

[34] Donnelly H, Ross E, Xiao Y, Hermantara R, Taqi AF, Doherty-Boyd WS, et al. Bioengineered niches that recreate physiological extracellular matrix organisation to support long-term haematopoietic stem cells. Nat Commun 2024;15. 10.1038/s41467-024-50054-0.

[35] Klein G. The extracellular matrix of the hematopoietic microenvironment. Experientia 1995;51:914–26. 10.1007/BF01921741.

[36] Bieniek M, Llopis-Hernandez V, Douglas K, Salmeron-Sanchez M, Lorenz C. Minor chemistry changes alter surface hydration to control fibronectin adsorption and assembly into nanofibrils. Adv Theory Simul 2019;1900169:1–13. 10.1002/adts.201900169.

[37] Alba-Perez A, Jayawarna V, Childs PG, Dalby MJ, Salmeron-Sanchez M. Plasma polymerised nanoscale coatings of controlled thickness for efficient solid-phase presentation of growth factors. Materials Science and Engineering C 2020;113. 10.1016/j.msec.2020.110966.

[38] Donnelly H, Ross E, Xiao Y, Hermantara R, Taqi AF, Doherty-Boyd WS, et al. Bioengineered niches that recreate physiological extracellular matrix organisation to support long-term haematopoietic stem cells. Nat Commun 2024;15. 10.1038/s41467-024-50054-0.

[39] Donnelly H, Sprott MR, Poudel A, Campsie P, Childs P, Reid S, et al. Surface-Modified Piezoelectric Copolymer Poly(vinylidene fluoride-trifluoroethylene) Supporting Physiological Extracellular Matrixes to Enhance Mesenchymal Stem Cell Adhesion for Nanoscale Mechanical Stimulation. ACS Appl Mater Interfaces 2023;15:50652–62. 10.1021/acsami.3c05128.

[40] Xiao Y, Donnelly H, Sprott M, Luo J, Jayawarna V, Lemgruber L, et al. Material-driven fibronectin and vitronectin assembly enhances BMP-2 presentation and osteogenesis. Mater Today Bio 2022;16. 10.1016/j.mtbio.2022.100367.

[41] Salmerón-Sánchez M, Dalby MJ. Synergistic growth factor microenvironments. Chem Commun 2016;52:13327–36. 10.1039/C6CC06888J.

[42] Chen X, Hughes R, Mullin N, Hawkins RJ, Holen I, Brown NJ, et al. Mechanical Heterogeneity in the Bone Microenvironment as Characterized by Atomic Force Microscopy. Biophys J 2020;119:502–13. 10.1016/j.bpj.2020.06.026.

[43] Jansen LA, Birch NP, Schiffman JD, Crosby AJ, Peyton SR. Mechanics of intact bone marrow. J Mech Behav Biomed Mater 2015;50:299–307. 10.1016/j.jmbbm.2015.06.023.

[44] Ligorio C, Zhou M, Wychowaniec JK, Zhu X, Bartlam C, Miller AF, et al. Graphene oxide containing self-assembling peptide hybrid hydrogels as a potential 3D injectable cell delivery platform for intervertebral disc repair applications. Acta Biomater 2019;92:92–103. 10.1016/j.actbio.2019.05.004.

[45] Simonson T, Brooks CL. Charge screening and the dieletric constant of proteins: Insights from molecular dynamics. J Am Chem Soc 1996;118:8452–8. 10.1021/ja960884f.

[46] Coutu DL, Kokkaliaris KD, Kunz L, Schroeder T. Three-dimensional map of nonhematopoietic bone and bone-marrow cells and molecules. Nat Biotechnol 2017;35:1202–10. 10.1038/nbt.4006.

[47] Pinho S, Frenette PS. Haematopoietic stem cell activity and interactions with the niche. Nat Rev Mol Cell Biol 2019;20:303–20. 10.1038/s41580-019-0103-9.

[48] Chen X, Hughes R, Mullin N, Hawkins RJ, Holen I, Brown NJ, et al. Mechanical Heterogeneity in the Bone Microenvironment as Characterized by Atomic Force Microscopy. Biophys J 2020;119:502–13. 10.1016/j.bpj.2020.06.026.

[49] Jansen LE, Birch NP, Schiffman JD, Crosby AJ, Peyton SR. Mechanics of intact bone marrow. J Mech Behav Biomed Mater 2015;50:299–307. 10.1016/j.jmbbm.2015.06.023.

[50] Pinho S, Lacombe J, Hanoun M, Mizoguchi T, Bruns I, Kunisaki Y, et al. PDGFR A and CD51 mark human stem cells capable of hematopoietic progenitor cell expansion. Journal of Experimental Medicine 2013;210:1351–67. 10.1084/jem.20122252.

[51] Kunisaki Y, Bruns I, Scheiermann C, Ahmed J, Pinho S, Zhang D, et al. Arteriolar niches maintain haematopoietic stem cell quiescence. Nature 2013;502:637–43. 10.1038/nature12612.

[52] Bandyopadhyay S, Duffy MP, Ahn KJ, Sussman JH, Pang M, Smith D, et al. Mapping the cellular biogeography of human bone marrow niches using single-cell transcriptomics and proteomic imaging. Cell 2024;187:3120–3140.e29. 10.1016/j.cell.2024.04.013.

[53] Asada N, Kunisaki Y, Pierce H, Wang Z, Fernandez NF, Birbrair A, et al. Differential cytokine contributions of perivascular haematopoietic stem cell niches. Nat Cell Biol 2017. 10.1038/ncb3475.

[54] Ding L, Morrison SJ. Haematopoietic stem cells and early lymphoid progenitors occupy distinct bone marrow niches. Nature 2013;495:231–5. 10.1038/nature11885.

[55] Engler AJ, Sen S, Sweeney HL, Discher DE. Matrix Elasticity Directs Stem Cell Lineage Specification. Cell 2006;126:677–89. 10.1016/j.cell.2006.06.044.

[56] Sugiyama T, Kohara H, Noda M, Nagasawa T. Maintenance of the Hematopoietic Stem Cell Pool by CXCL12-CXCR4 Chemokine Signaling in Bone Marrow Stromal Cell Niches. Immunity 2006;25:977–88. 10.1016/j.immuni.2006.10.016.

[57] Dorrell C, Gan OI, Pereira DS, Hawley RG, Dick JE. Expansion of human cord blood CD34+CD38−cells in ex vivo culture during retroviral transduction without a corresponding increase in SCID repopulating cell (SRC) frequency: dissociation of SRC phenotype and function. Blood 2000;95:102–10. 10.1182/blood.V95.1.102.

[58] Wisniewski D, Affer M, Willshire J, Clarkson B. Further phenotypic characterization of the primitive lineage− CD34+CD38−CD90+CD45RA− hematopoietic stem cell/progenitor cell sub-population isolated from cord blood, mobilized peripheral blood and patients with chronic myelogenous leukemia. Blood Cancer J 2011;1:e36– e36. 10.1038/bcj.2011.35.

[59] Van Galen P, Kreso A, Wienholds E, Laurenti E, Eppert K, Lechman ER, et al. Reduced lymphoid lineage priming promotes human hematopoietic stem cell expansion. Cell Stem Cell 2014;14:94–106. 10.1016/j.stem.2013.11.021.

[60] Campa CC, Weisbach NR, Santinha AJ, Incarnato D, Platt RJ. Multiplexed genome engineering by Cas12a and CRISPR arrays encoded on single transcripts. Nat Methods 2019;16:887–93. 10.1038/s41592-019-0508-6.

[61] Cong L, Ran FA, Cox D, Lin S, Barretto R, Hsu PD, et al. Multiplex Genome Engineering Using CRISPR/Cas Systems Le. Science 2013;339:819–23. 10.1126/science.1231143.Multiplex.

[62] Borot F, Wang H, Ma Y, Jafarov T, Raza A, Ali AM, et al. Gene-edited stem cells enable CD33-directed immune therapy for myeloid malignancies. Proc Natl Acad Sci U S A 2019;116:11978–87. 10.1073/pnas.1819992116.

[63] Fioretti S, Matson CA, Rosenberg KM, Singh NJ. Host B cells escape CAR-T immunotherapy by reversible downregulation of CD19. Cancer Immunology, Immunotherapy 2023;72:257–64. 10.1007/s00262-022-03231-3.

[64] Llopis-Hernández V, Cantini M, González-García C, Cheng ZA, Yang J, Tsimbouri PM, et al. Material-driven fibronectin assembly for high-efficiency presentation of growth factors. Sci Adv 2016;2:1–11. 10.1126/sciadv.1600188.

[65] Kaplan FS, Xu M, Seemann P, Connor JM, Glaser DL, Carroll L, et al. Classic and atypical fibrodysplasia ossificans progressiva (FOP) phenotypes are caused by mutations in the bone morphogenetic protein (BMP) type I receptor ACVR1. Hum Mutat 2009;30:379–90. 10.1002/humu.20868.

[66] Brazil DP, Church RH, Surae S, Godson C, Martin F. BMP signalling: Agony and antagony in the family. Trends Cell Biol 2015;25:249–64. 10.1016/j.tcb.2014.12.004.

[67] Xiao Y, Donnelly H, Sprott M, Luo J, Jayawarna V, Lemgruber L, et al. Material-driven fibronectin and vitronectin assembly enhances BMP-2 presentation and osteogenesis. Mater Today Bio 2022;16:100367. 10.1016/j.mtbio.2022.100367.

[68] Chaudhuri O, Cooper-White J, Janmey PA, Mooney DJ, Shenoy VB. Effects of extracellular matrix viscoelasticity on cellular behaviour. Nature 2020;584:535–46. 10.1038/s41586-020-2612-2.

[69] Méndez-Ferrer S, Michurina T V., Ferraro F, Mazloom AR, MacArthur BD, Lira SA, et al. Mesenchymal and haematopoietic stem cells form a unique bone marrow niche. Nature 2010;466:829–34. 10.1038/nature09262.

[70] Pinho S, Lacombe J, Hanoun M, Mizoguchi T, Bruns I, Kunisaki Y, et al. PDGFRα and CD51 mark human Nestin+ sphere-forming mesenchymal stem cells capable of hematopoietic progenitor cell expansion. Journal of Experimental Medicine 2013;210:1351–67. 10.1084/jem.20122252.

[71] King PJS, Giovanna Lizio M, Booth A, Collins RF, Gough JE, Miller AF, et al. A modular self-assembly approach to functionalised β-sheet peptide hydrogel biomaterials. Soft Matter 2016;12:1915–23. 10.1039/c5sm02039e.

[72] Rodriguez-Fraticelli AE, Wolock SL, Weinreb CS, Panero R, Patel SH, Jankovic M, et al. Clonal analysis of lineage fate in native haematopoiesis. Nature 2018;553:212–6. 10.1038/nature25168.

[73] Kim MY, Yu KR, Kenderian SS, Ruella M, Chen S, Shin TH, et al. Genetic Inactivation of CD33 in Hematopoietic Stem Cells to Enable CAR T Cell Immunotherapy for Acute Myeloid Leukemia. Cell 2018;173:1439–1453.e19. 10.1016/j.cell.2018.05.013.

[74] Frangoul H, Altshuler D, Cappellini MD, Chen Y-S, Domm J, Eustace BK, et al. CRISPR-Cas9 Gene Editing for Sickle Cell Disease and β-Thalassemia. New England Journal of Medicine 2021;384:252–60. 10.1056/nejmoa2031054.

[75] Tambaro FP, Singh H, Jones E, Rytting M, Mahadeo KM, Thompson P, et al. Autologous CD33-CAR-T cells for treatment of relapsed/refractory acute myelogenous leukemia. Leukemia 2021;35:3282–6. 10.1038/s41375-021-01232-2.

[76] Mishra A, Maiti R, Mohan P, Gupta P. Antigen loss following CAR-T cell therapy: Mechanisms, implications, and potential solutions. Eur J Haematol 2024;112:211–22. 10.1111/ejh.14101.

[77] Gardner R, Wu D, Cherian S, Fang M, Hanafi LA, Finney O, et al. Acquisition of a CD19-negative myeloid phenotype allows immune escape of MLL-rearranged B-ALL from CD19 CAR-T-cell therapy. Blood 2016;127:2406–10. 10.1182/blood-2015-08-665547.

[78] Dickinson AM, Norden J, Li S, Hromadnikova I, Schmid C, Schmetzer H, et al. Graft-versus-Leukemia Effect Following Hematopoietic Stem Cell Transplantation for Leukemia. Front Immunol 2017;8. 10.3389/fimmu.2017.00496.

[79] Kouro T, Himuro H, Sasada T. Exhaustion of CAR T cells: potential causes and solutions. J Transl Med 2022;20:1–10. 10.1186/s12967-022-03442-3.

[80] Alba-Perez A, Jayawarna V, Childs PG, Dalby MJ, Salmeron-Sanchez M. Plasma polymerised nanoscale coatings of controlled thickness for efficient solid-phase presentation of growth factors. Materials Science and Engineering C 2020;113:110966. 10.1016/j.msec.2020.110966.

[81] Cantini M, Rico P, Moratal D, Salmer On-S Anchez M. Controlled wettability, same chemistry: biological activity of plasma-polymerized coatings. Soft Matter8 2012;8:5575–84. 10.1039/c2sm25413a.

[82] Brinkman EK, Chen T, Amendola M, Van Steensel B. Easy quantitative assessment of genome editing by sequence trace decomposition. Nucleic Acids Res 2014;42:1–8. 10.1093/nar/gku936.

