## Supplementary figures for "Synthetic peptide hydrogels as a model of the bone marrow niche demonstrate efficacy of a combined CRISPR-CAR T-cell therapy for acute myeloid leukaemia"

### Supplementary figure legends

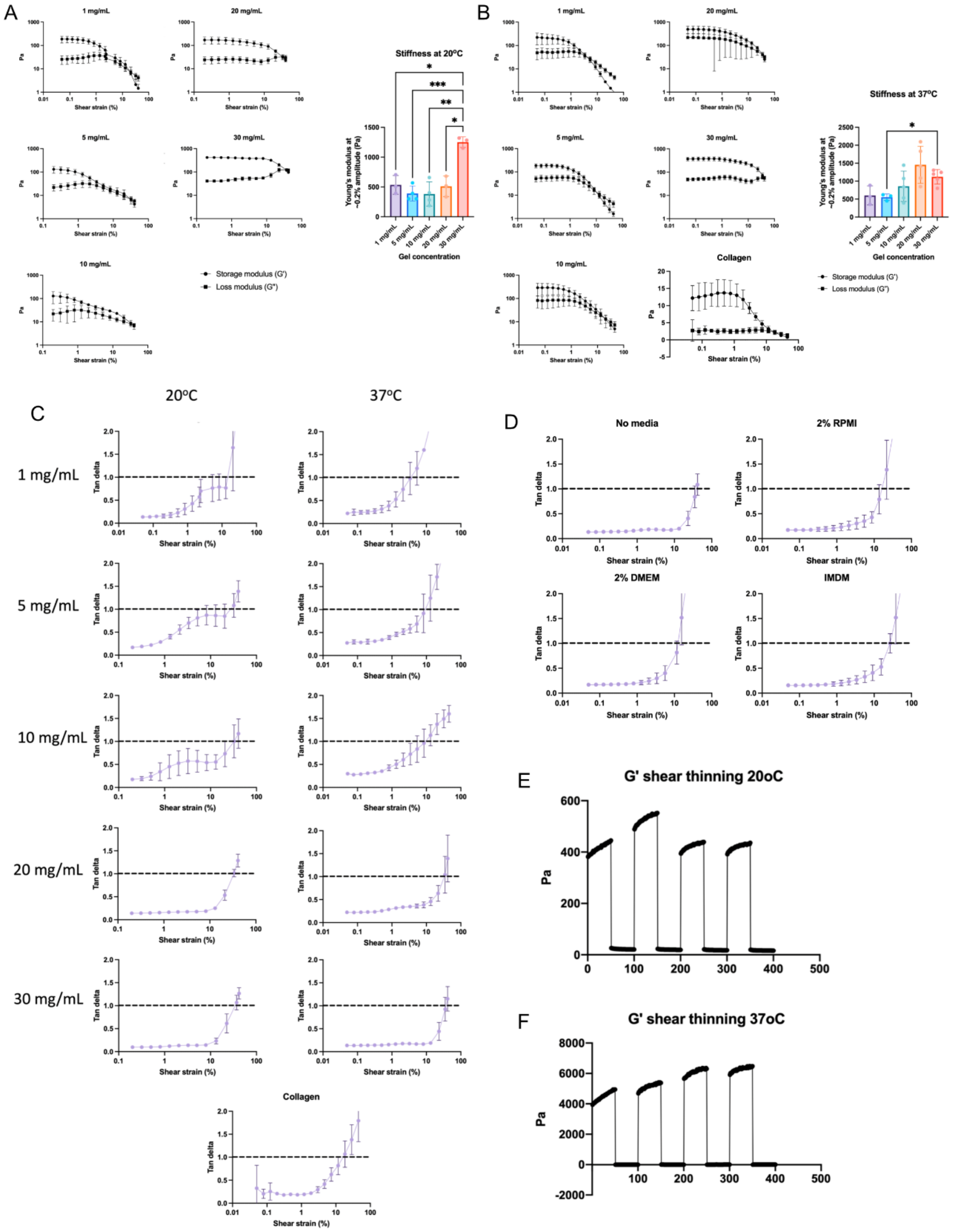


***Supplementary figure 1. Further characterisation of gels' mechanical properties. A-B*** *Effect of gel dilution on G', G" and Young's modulus of PeptiGel at* ***A*** *20°C and* ***B*** *37°C, assessed with rheological analysis. Collagen was also assessed at 37°C.* ***C*** *Tan delta of various PeptiGel dilutions at 20°C or 37°C, and collagen gel at 37°C.* ***D*** *Effect of media on PeptiGel tan delta.* ***E*** *Stress recovery of PeptiGels at 20°C or 37°C reveals shear thinning properties. Graphs show mean ± SD. Statistics for* ***A*** *and* ***B*** *by one-way Brown-Forsyth and Welch ANOVA followed by Dunnett T3 multiple comparison test. *$p < 0.05$, **$p < 0.01$, ***$p < 0.001$ and ****$p < 0.0001$. Non-significant not shown.* ***A-D*** *n=3 experimental replicates,* ***E-F*** *n=1 experimental replicate.*

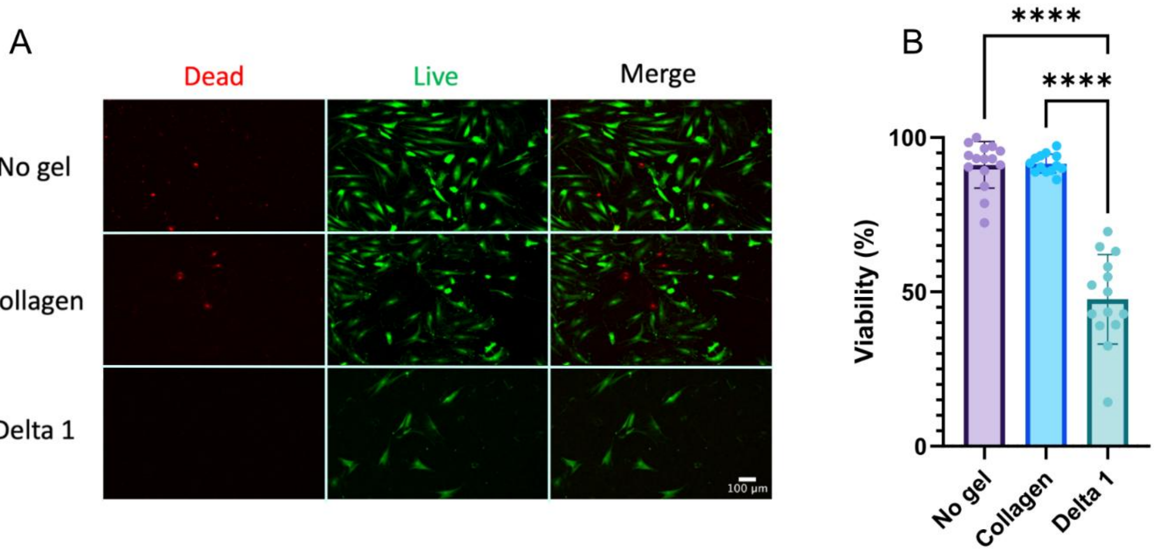


***Supplementary figure 2. Viability analysis of MSCs in various conditions. A*** *Representative images of viability assay of MSCs in various gel conditions.* ***B*** *Viability of MSCs in various gel conditions. For representative images, scale bars are 100 µm, red = dead, ethidium homodimer-stained cells, green = live, calcein-stained cells. Graphs show mean ± SD. Statistics for* ***B*** *by Kruskal-Wallis followed by Dunn's multiple comparison test. *p < 0.05, **p < 0.01, ***p < 0.001 and ****p < 0.0001, non-significant not shown. n=3 experimental replicates, 3 or 4 photos taken per replicate and individually assessed for % viability.*

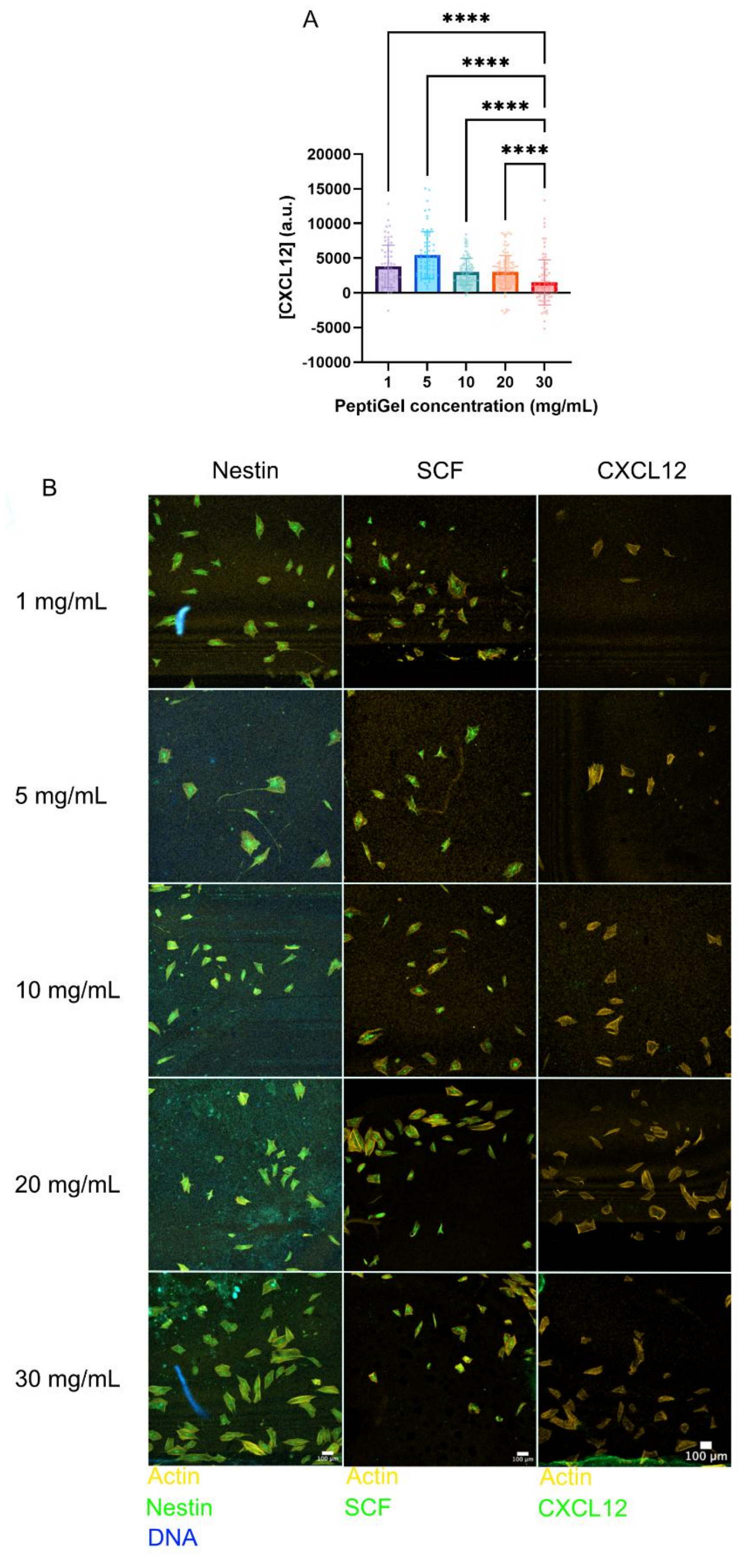


***Supplementary figure 3. Effect of PeptiGel concentration on MSC phenotype. A*** *Effect of gel dilution on CXCL12, assessed by IF microscopy.* ***B*** *Representative IF microscopy images of MSCs under 1-30 mg/mL PeptiGel. Scale bar is 100 μm, yellow = in all images actin, green = from left to right columns: nestin, SCF, CXCL12, blue = DNA in first column. Graphs show mean ± SD. Statistics for* ***A*** *by Kruskal-Wallis followed by Dunn's multiple comparison test. *p < 0.05, **p < 0.01, ***p < 0.001 and ****p < 0.0001, non-significant not shown. n=3 experimental replicates of ~10 cells that were individually analysed.*

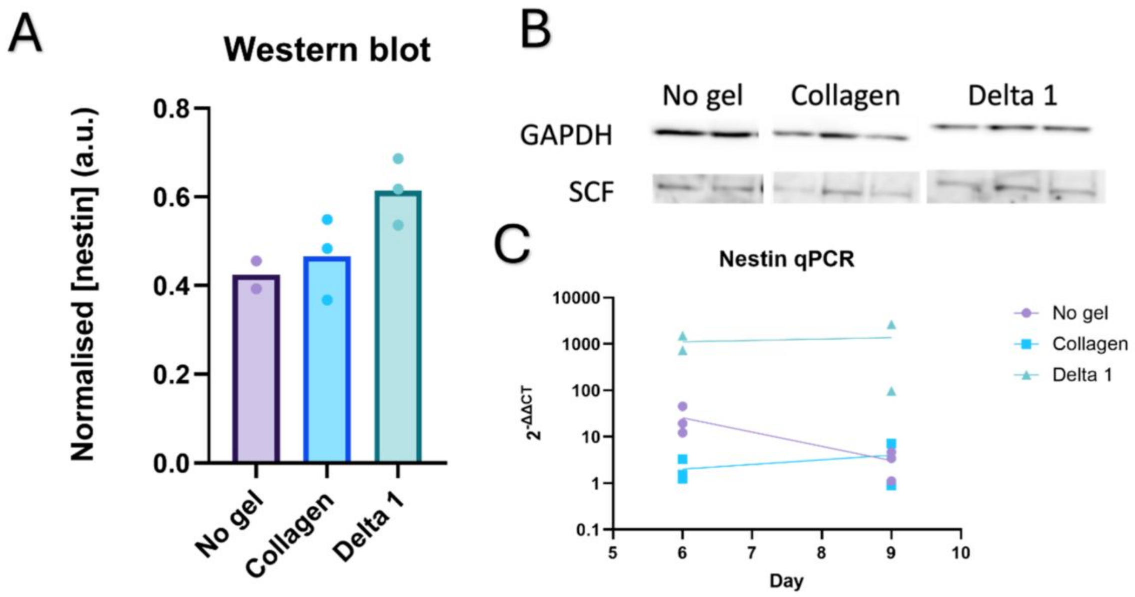


***Supplementary figure 4. Western blot and qPCR analysis of effect of gels on MSCs' phenotype. A*** *Western blot analysis of MSCs' SCF expression in no gel, collagen and PeptiGel conditions.* ***B*** *Representative images of SCF and GAPDH western blots.* ***C*** *qPCR analysis of MSCs' nestin expression in PEA-FN-BMP2 +no gel/collagen/PeptiGel conditions, normalised against GAPDH expression.* ***A*** *n=2 for no gel condition, and n=3 for collagen and PeptiGel conditions.* ***B*** *n=2 for collagen day 9 and for PeptiGel days 6 and 9, n=3 for other conditions and timepoints.*

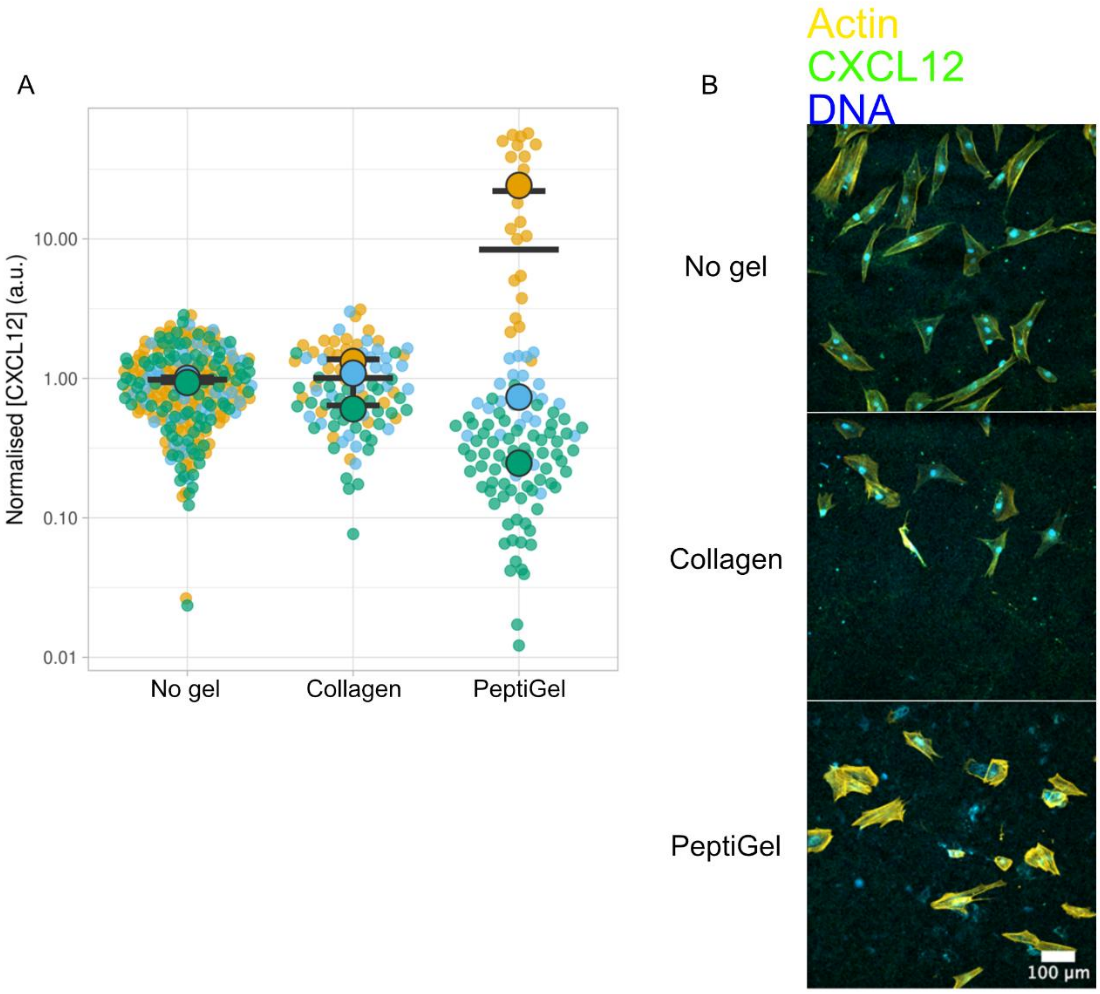


***Supplementary figure 5. Effect of gels on MSCs' CXCL12 expression. A*** *CXCL12 expression of MSCs in no gel, collagen and PeptiGel conditions, assessed by IF microscopy.* ***B*** *Representative images of MSCs. Scale bar is 100 µm, yellow = actin, green = CXCL12, blue = DNA. Graph shows mean ± SD. n=3 biological replicates, each consisting of 4 experimental replicates.*

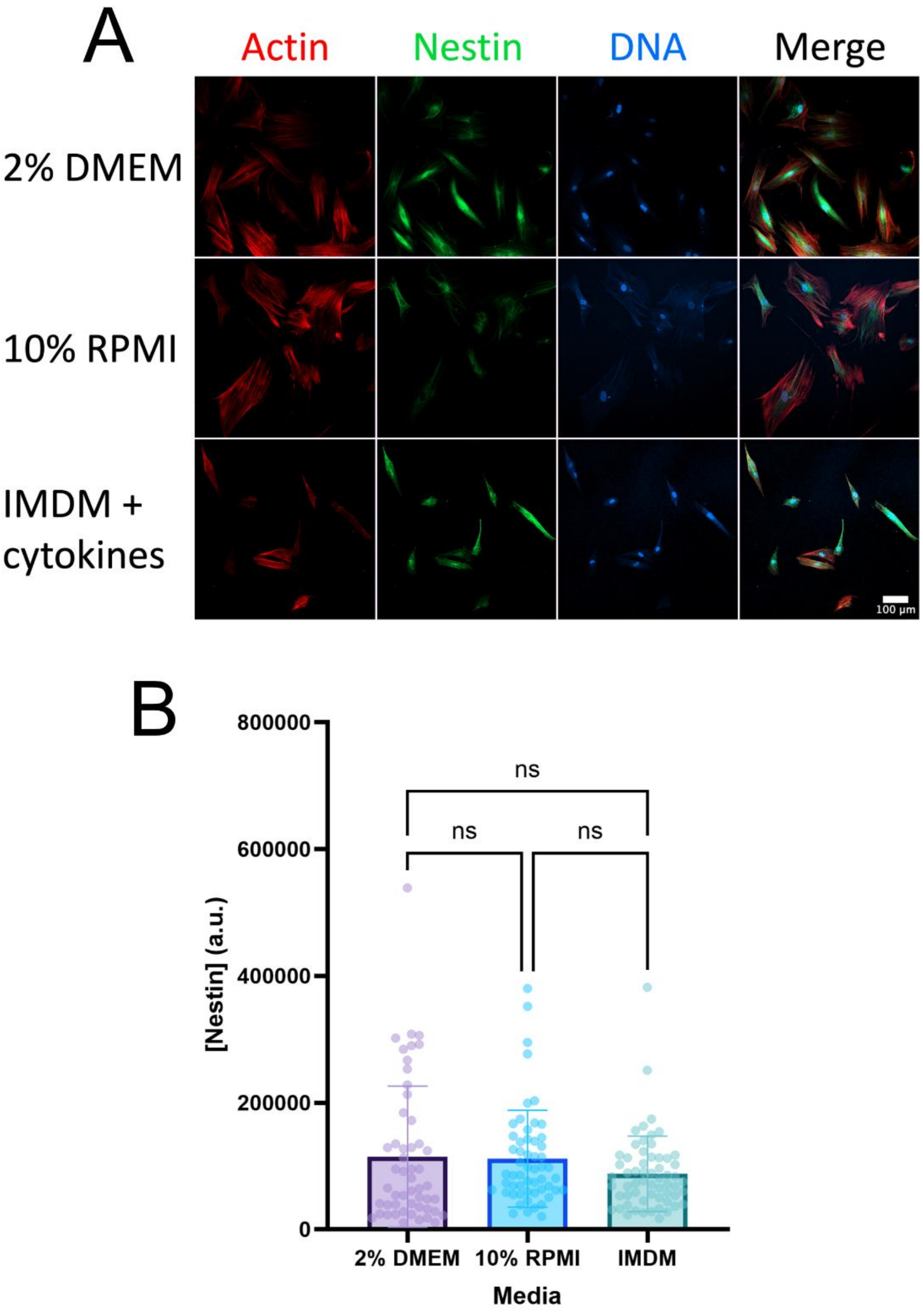


***Supplementary figure 6. Effect of media on MSCs' phenotype in model systems. A** Representative image of MSCs cultured with various media formulae. Scale bar is 100 μm, red = actin, green = nestin, blue = DNA. **B** Effect of media on MSCs' nestin expression. Graph shows mean ± SD. Statistics by Kruskal-Wallis followed by Dunn's multiple comparison test. *p < 0.05, **p < 0.01, ***p < 0.001 and ****p < 0.0001, non-significant shown as ns. n=3, with ~10 cells individually analysed per repeat.*

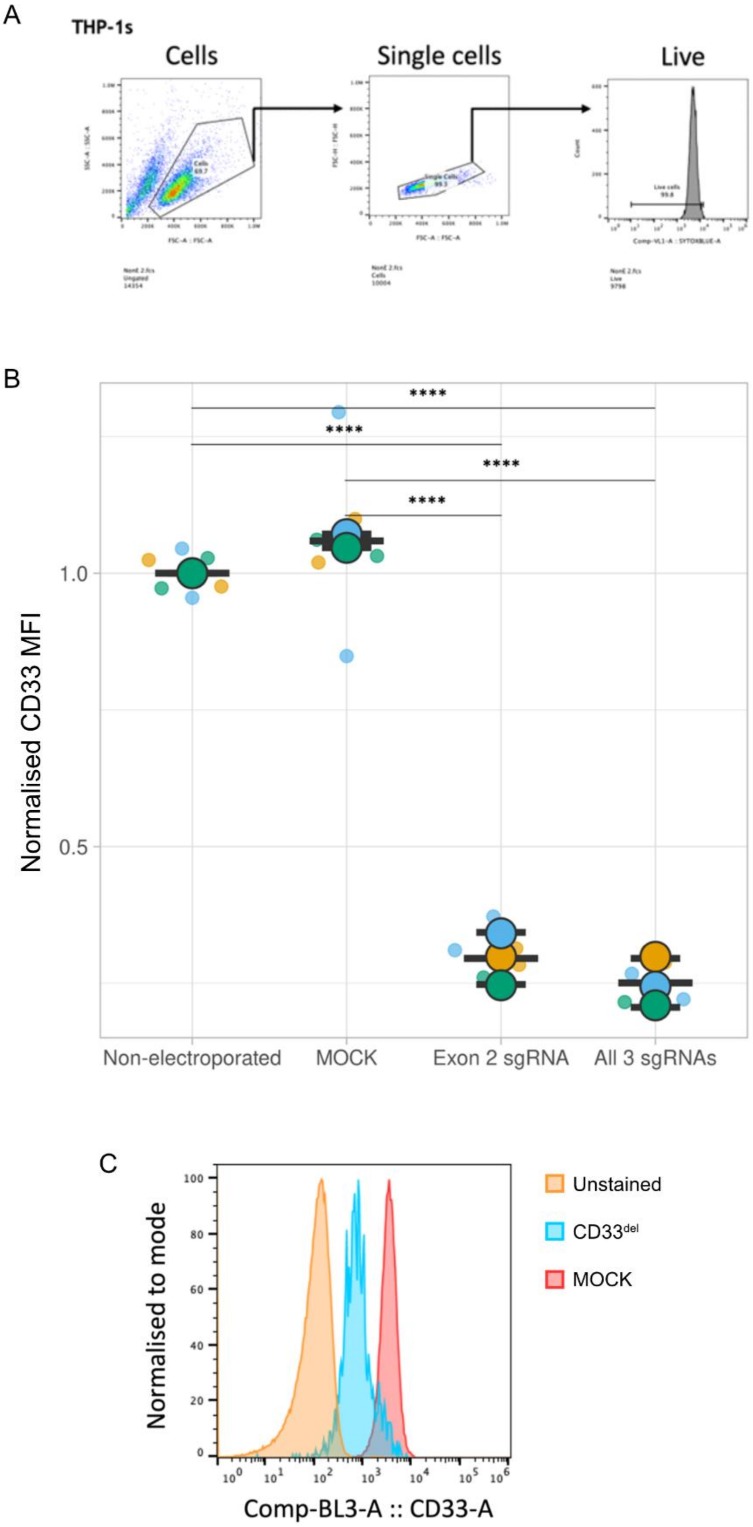


***Supplementary figure 7. CD33 efficiently disrupted in THP-1s. A** Flow cytometry gating strategy for assessing THP-1s' CD33 concentration on cells' surfaces. **B** CD33 expression (MFI) assessed by flow cytometry of THP-1s that were non-electroporated, MOCK treated, edited with a single sgRNA targeting CD33 exon 2, or edited with a cocktail of all three sgRNAs. **C** Representative flow cytometry fluorescence histogram of THP-1s that were unstained, MOCK treated and edited with a cocktail of all three sgRNAs ($CD33^{del}$)**.** Graph shows mean ± SD. Statistics by one-way ANOVA followed by Tukey multiple comparison test. *$p < 0.05$, **$p < 0.01$, ***$p < 0.001$ and ****$p < 0.0001$, non-significant not shown. n=3 independent replicates using different cultures of the THP-1 cell line. Each replicate consisted of 2 experimental replicates.*

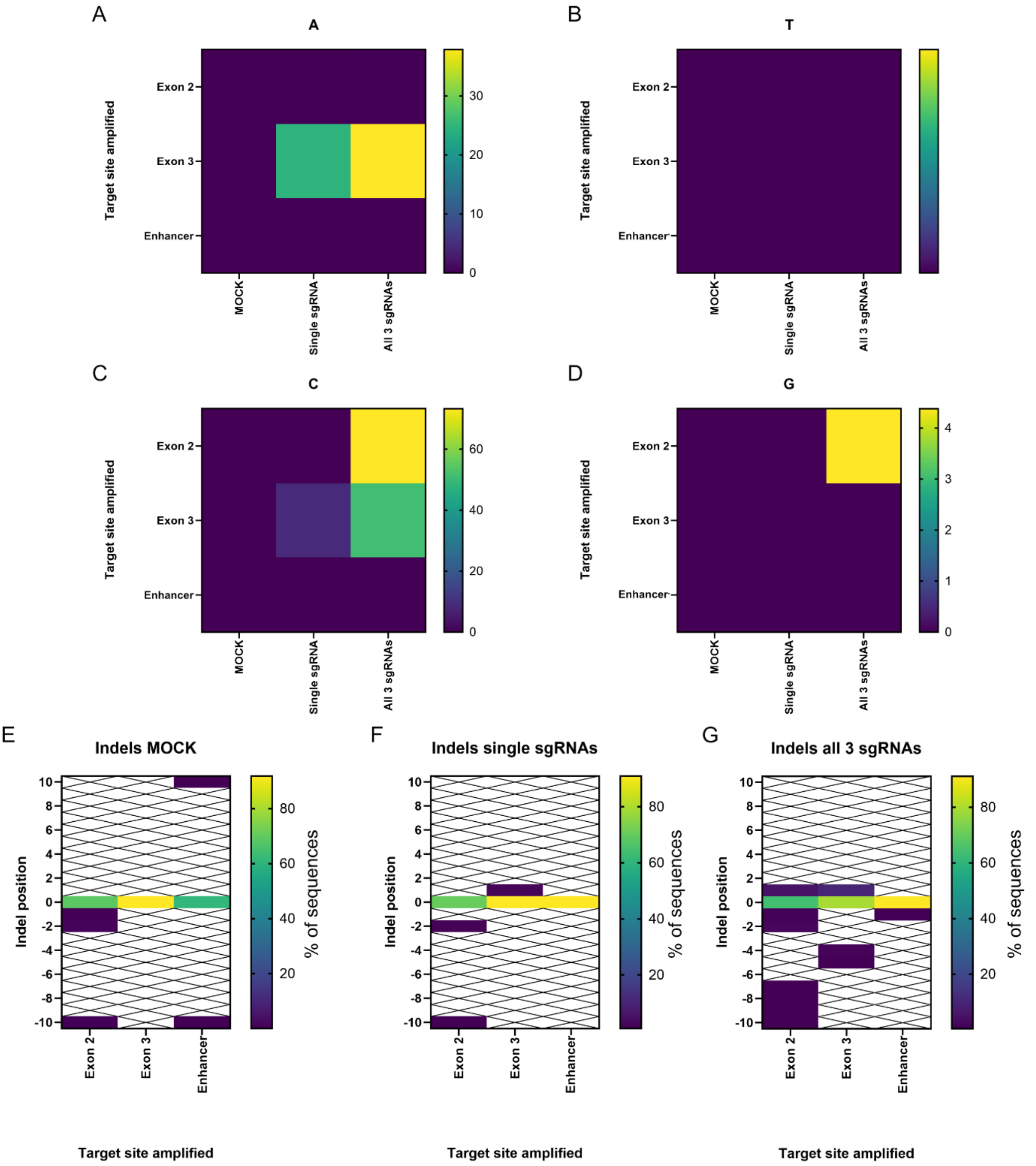


***Supplementary figure 8. TIDE characterisation of CD33 disruption in HSCs. A-D*** *Characterisation of +1 insertions when no sgRNA was used (MOCK), a single sgRNA targeting the analysed target site (exon 2 in CD33, exon 3 in CD33, or an upstream CD33 enhancer) was used, or a cocktail of sgRNAs targeting sites in exon 2, exon 3, and an upstream promoter of CD33 were used. Each base is represented by a dedicated heat map:* ***A*** *adenine,* ***B*** *thymine,* ***C*** *cytosine,* ***D*** *guanine.* ***E-G*** *Indel characterisation of* ***E*** *MOCK conditions,* ***F*** *single sgRNA conditions, and* ***G*** *cocktail of three sgRNA conditions, showing the location and percentage frequency of indels relative to the expected cut site (represented as 0). A higher percentage of 0 values indicates greater alignment of the control and test sequences, implying lower mutagenesis. White boxes with a black cross indicate that no statistically significant indels were detected at the corresponding site for the corresponding condition. Heat maps show mean. n=3 experimental replicates.*
